# Capacitance Sensor Array for Lab-on-CMOS Applications using a Passive RFID Interface

**DOI:** 10.64898/2026.02.05.704137

**Authors:** Kai-Chun Lin, Marc Dandin

**Affiliations:** Department of Electrical and Computer Engineering, Carnegie Mellon University, Pittsburgh, PA 15213 USA

**Keywords:** Biosensor, capacitance sensor, CMOS, lab-on-a-chip, lab-on-CMOS, near-field, on-chip antenna, radio frequency, RF front-end, RFID, passive RFID, UHF

## Abstract

We report a 0.18 *µ*m CMOS lab-on-a-chip system that monolithically integrates a *passive* radio frequency identification (RFID) interface and an 8 × 8 array of capacitance sensors configured for measuring the capacitance change resulting from an overlying biological specimen. This lab-on-CMOS platform is designed to operate wirelessly, first in a *harvesting* mode in which on-chip power is generated via the inductive coupling of an on-chip antenna to an external antenna, and second, in a *sense-and-transmit* mode where the capacitance sensor array is scanned and the measured data are transmitted to the external antenna using the same on-chip antenna. This paper presents characterization results of the passive RFID interface and of the sensor core, the latter utilizing several test analytes. The proposed system will facilitate the integration and packaging of a large number of chips in wet environments, paving the way for the inclusion of lab-on-CMOS technology in standard bio-analytical lab practice.

## I. Introduction

**C**APACITANCE sensors are used in biomedical research to quantify the interactions of biological specimen with a sensing electrode. The real-time monitoring of these interactions can yield important cues that inform on biological activity. For instance, capacitance sensors can be used to characterize the morphology and proliferation of cancer cells, the formation of bacterial biofilms, the activation of immune cells by nanoparticles, the onset of viral exposure in cells, and DNA hybridization [1]–[5].

When implemented in complementary metal-oxide semiconductor (CMOS) processes, capacitance sensors can be integrated with cell culture apparatuses such as microfluidic channels or cell culture wells to form a compact bio-analytical microsystem termed a “lab-on-CMOS.” In this paradigm, biosensing functions are realized directly on the CMOS chip’s surface [6], [7].

There are significant challenges to this integration process [8]–[10]. For example, creating a wet environment for cells to thrive while maintaining the CMOS chip’s electrical functions is not trivial. Specifically, while microfluidic networks readily provide a means to deliver samples at sensing locations, the liquid media they contain must not come in contact with the wirebonds that connect the chip to its package. Even when such contact is prevented by encapsulating the wirebonds using a dielectric material, during longtime measurements, the ionic liquid media may diffuse into the encapsulant. This may create electrical conduction paths between otherwise electrically isolated wires, which ultimately leads to electrical failure [11].

One approach to mitigating these problems has been to embed the chip in a recessed carrier and providing planar metal interconnects to an outer pad frame from which wirebonds can be made, far away from the sensing area (Fig. 2) [12]–[15]. Such integrative packaging processes have been successful in providing prototypes that can last long enough to carry out experiments. However, in each case, extensive post-CMOS fabrication was required. Therefore, this option may be cost-prohibitive.

Furthermore, the yield of the integration process may be low due to the difficulties in bridging the step between the chip carrier and the chip with a continuous metal layer [8]. Lastly, this approach tends to be difficult to scale in applications where many chips are to be integrated within the same microsystem, owing to the increased complexity that arises when routing I/Os from each chip to an outer redistribution pad frame.

Thus, to overcome the aforementioned limitations, we propose a novel RF-enabled lab-on-CMOS (RFLoC) capacitance sensor that includes an integrated passive RFID interface, as shown in Fig. 3. The device requires no battery as it can harness energy from the electromagnetic field established by an external antenna to generate on-chip power, sense, and report data wirelessly. In sum, we propose an approach that will reduce the complexity of the packaging process by obviating the need for wirebonds or post-CMOS microfabrication.

Our approach will also facilitate new applications wherein multiple devices are integrated on the same platform to provide massively-parallel assays. In such applications, a plurality of RFID-enabled sensor chips may be accessed using a multi-access RFID protocol designed to avoid data collisions [16]– [18].

This paper presents characterization results from a prototype RFLoC chip designed according to the proposed passive RFID framework. The chip was implemented in a 0.18 *µ*m twin-well, 2-poly, and 6-metal commercial CMOS process. At the core, it includes a cpacitance-to-frequency sensor, leveraging our previous work on capacitance-sensing lab-on-CMOS microsystems [19]–[23]. We discuss the system’s architecture, and we present measurement results from several analytes using the chip’s sensor core. Characterization results of the chip’s wireless power transfer efficiency and data uplink performance are also reported for several test conditions.

## II. System Design

### A. System Architecture & CMOS Implementation

A block diagram of the proposed RFLoC concept is shown in Fig. 4. Generally, the architecture includes a sensor core formed by a capacitance-sensing pixel array, RF front-end circuitry for power management and wireless communication, and data processing circuits. The sensor core consists of an 8×8 array of capacitance-to-frequency biosensors (discussed at length *infra*), an address decoder, and an output multiplexer. The data processing block includes a 24-bit counter, a parallel-in-series-out (PISO) interface circuit, a current bias generation circuit, voltage reference circuits, and a data modulation circuit.

The planar OCA interfaces with the power management system via a programmable resonant frequency tuning tank and via a transmitter configured to send measurement data out wirelessly. The RF front-end further includes a shunt regulator to regulate the supply voltage. The chip is operable with a 900 *∼*920 MHz carrier signal, *i*.*e*., with a signal in the ISM band of the UHF spectrum. The carrier signal is generated via an external reader antenna that inductively couples with the OCA to deliver wireless power. The external reader antenna is also configured to receive sensor output data streams sent by the on-chip transmitter. The tuning tank is configured to tune the resonant frequency of the RFLoC’s OCA to match the carrier frequency and achieve maximum wireless power transfer.

During operation, the boost rectifier converts the induced RF signal to a DC voltage *V*_*DD*_ which is regulated by the shunt regulator to a desired level in order to power the various circuitries on the chip. Following this harvesting mode, the chip enters a sense-and-transmit mode wherein the pixel array reports sensor data to the data processing circuits which modulate the data and ready them for subsequent transmission by the RF front-end.

Fig. 5 depicts the floor plan for an implementation of the RFLoC in a 0.18 *µ*m commercial CMOS process. We note that the prototype is designed for testability. In other words, I/O pads and scan chain hardware were included in order to test the system’s various components and sub-components independently, and to flexibly control sensor pixel addressing. In future implementations, an automatic pixel array scanning feature will be included, allowing the chip to function in a fully autonomous manner within the proposed wireless communication framework.

In the instant implementation, the OCA surrounds the 8×8 sensor array and the various other circuitries comprised in the chip. The timing diagram illustrating the chip’s operation is depicted in Fig. 6. First, the chip establishes a *V*_*DD*_ from the reader wirelessly. Second, a reset signal toggles to reset the internal circuits. Third, address information is programmed into the chip via the scan chain to activate the selected sensor pixel. The counter is also enabled to record and store the clock counts accordingly. Fourth, the counter stops when the specified measurement time has elapsed, and fixed pattern “preamble” and “appendix” bits are added at the beginning and at then of the sensor’s output data bits stored in the output counter.

Lastly, a PISO synchronization circuit generates an internal data-fetch pulse lasting at least one clock cycle to ensure that the PISO circuit fetches the counter data correctly. The PISO interface then converts the parallel data bits into a series output data stream and passes it to the transmitter. The data are transmitted to the external reader antenna.

### B. Pixel Architecture & Theory of Operation

The RFLoC’s biosensor pixels consist of *circuit-under-pad* oscillator-based capacitance-to-frequency sensors. Specifically, each sensor pixel in the array consists of a three-stage current-starved ring oscillator whose intermediate stage is connected to a metal-oxide-metal sensing electrode implemented with the top metal layer of the process. The circuit-under-pad design is achieved by inserting a ground metal shield between the ring oscillator circuit and the electrode in order to reduce noise coupling from the oscillator; this feature also serves to minimize pixel area, which facilitates placing the 8×8 array within the OCA’s inner loop.

When a cell under study adsorbs unto the chip’s passivation layer directly over the sensing electrode, as described for example in Fig. 1, the electrode’s effective capacitance changes, and it is this change that is correlated with cell activity at the bioelectronic interface. This effective capacitance is written in Equation 1 as a capacitance (*C*_*coupled*_) that couples to the oscillator as result of the cell being within the electrode’s vicinity. It includes the overlap capacitance *C*_*ov*_ of the electrode and the sensed capacitance *C*_*T*_ due to the cell as well as the deterministic parasitic capacitances of the CMOS stack. In our implementation, the electrode is formed using a number of interdigitated top metal fingers (Fig. 7) represented by the parameter *n* in Equation 1. Further, in Equation 1, *ϵ*_0_ is the dielectric constant of free space, *ϵ*_*r*_ the dielectric constant of the non-conductive region between the fingers, *A* the area of the plate formed by each finger, and *d* the distance between two fingers.

**Fig. 1:**
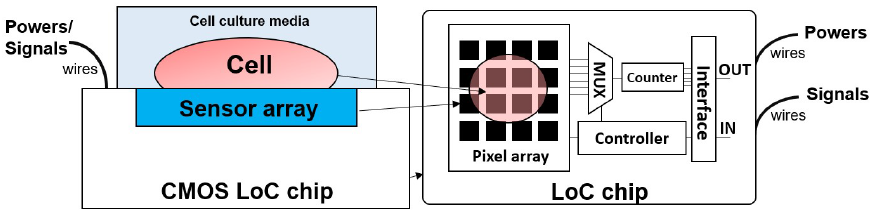
Cross-sectional (left) and top (right) views of a conventional capacitance-sensing lab-on-CMOS system. The CMOS chip includes an array of passivated electrodes connected to circuitry that measures cell-substrate coupling as a change in capacitance at the interface between the electrodes and the cells. The chip further includes readout circuits for routing data off-chip via wirebonds. Power and ground lines are also provided via wirebonds.

**Fig. 2:**
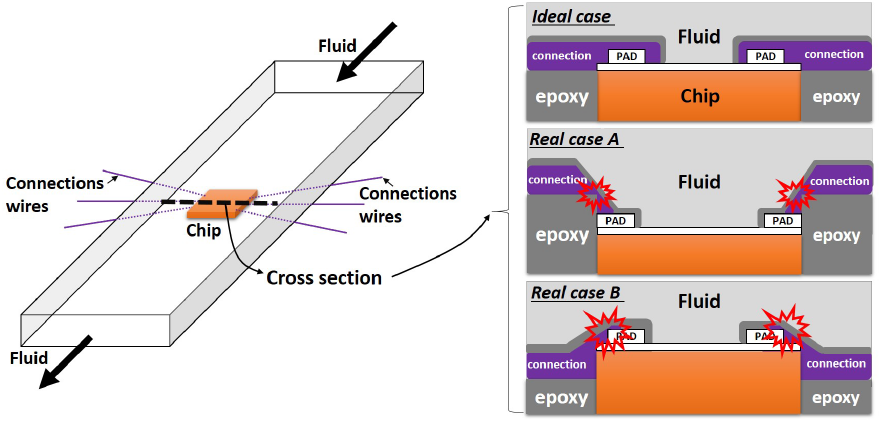
Left: microfluidics integration in a lab-on-CMOS system. The chip is placed inside a microchannel, and its input/output (I/O) pads are connected to an outer pad frame (not shown) using planar metal interconnects, which are subsequently passivated. Right: cross-sectional views showing an ideal device structure after integration, and two practical cases which highlight potential areas of electrical connection failure.

**Fig. 3:**
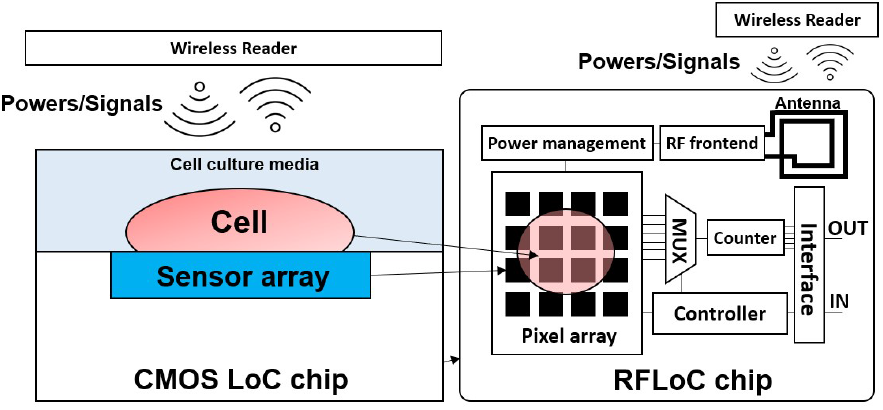
Cross-sectional (left) and top view (right) of the proposed RF-enabled lab-on-CMOS microsystem. The device features an RF-front-end, an on-chip antenna, on-board power management circuits, a sensor core, and readout circuits. Power is induced on the chip via the inductive coupling between the on-chip antenna and the antenna of an external wireless reader device.

**Fig. 4:**
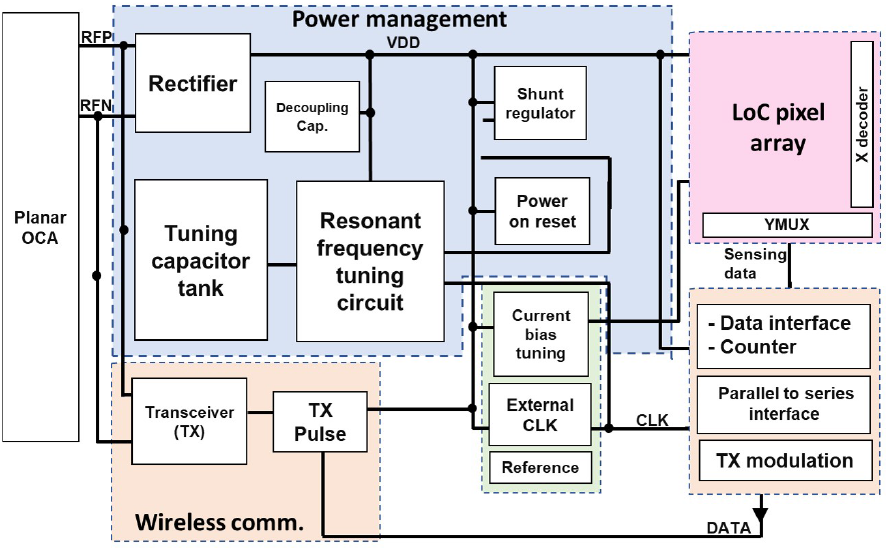
Block diagram of the proposed RF-enabled lab-on-CMOS (RFLoC) system capacitance-sensing system. The chip is designed to operate in the ISM sub-band of the UHF radio spectrum. It includes an on-chip antenna and monolithically-integrated circuitry for power management, data processing, biosensing, and wireless communication.

**Fig. 5:**
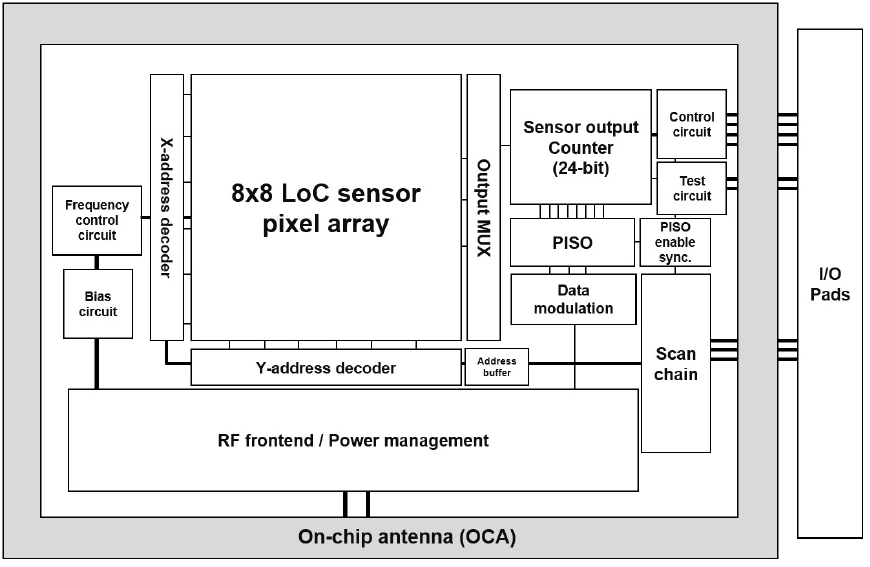
Floor plan of the RFLoC implemented in a 0.18 *µ*m twin-well, 2-poly, and 6-metal commercial CMOS process. The LoC sensor core and the chip’s other constitutive circuit blocks are surrounded by the OCA.The I/O pads are placed outside the OCA to provide independent testing functionality for the various chip sub-components.

**Fig. 6:**
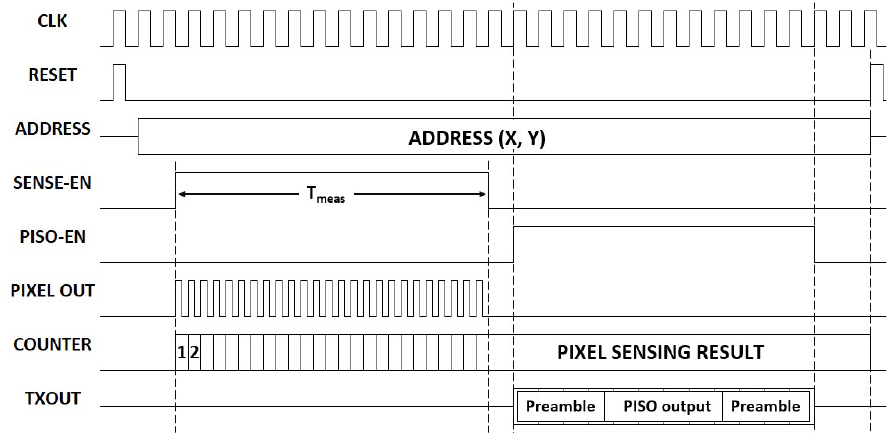
Timing diagram for the chip’s operation. The *V*_*DD*_ power supply is induced by coupling the external antenna to the OCA. The chip then resets all circuits, performs a sensing operation, and packages the data for transmission via the RF front-end.

**Fig. 7:**
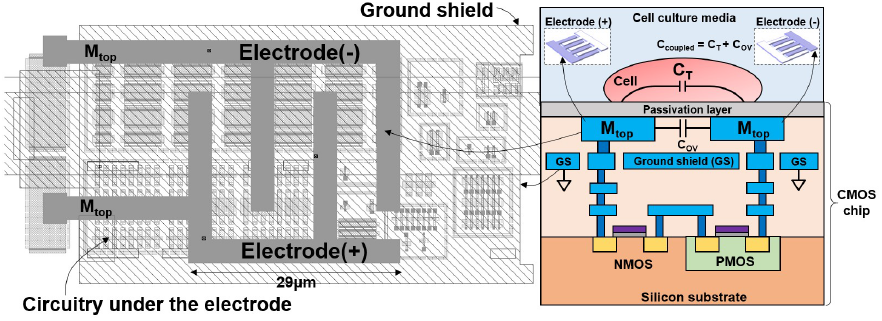
Layout view (left) and cross-sectional view (right) of the circuit-underpad pixel design. The electrode is constructed with the process’ top thick metal, and a ground plane is placed under the electrode to shield it from the noise originating from the underlying CMOS circuitry.

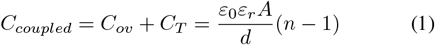

Fig. 8 shows the pixel circuit architecture. It consists of a three-stage current-starved ring oscillator, which can be enabled upon pixel selection. The base current at each stage of the oscillator is controllable via a programmable current source, *i*.*e*., the output frequency of the oscillator can be tuned by changing the current reference option codes P*<*3:0*>* and N*<*2:0*>* externally.

**Fig. 8:**
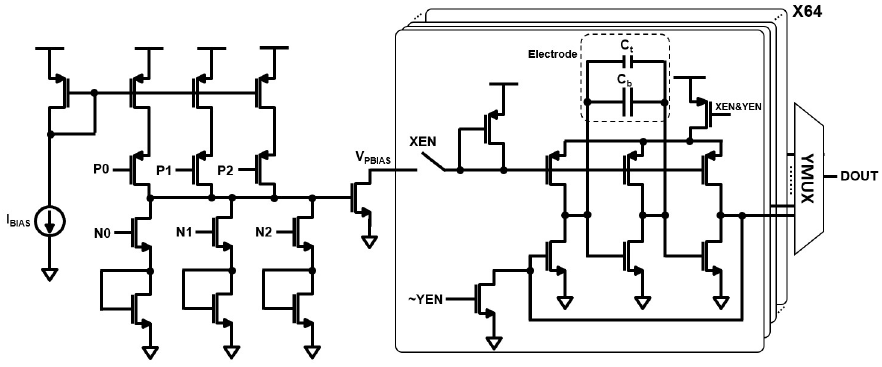
Circuit diagram of the 64-pixel capacitance sensor core. The array is interfaced with a global and controllable current reference. Each pixel has a 3-stage current-starved NMOS oscillator that is connected to an output multiplexer.

The three-stage current-starved ring oscillator acts as the sensing circuit when the two interdigitated electrode leads are connected to the input and output nodes of the oscillator’s middle stage. Any change in *C*_*coupled*_ alters the effective gate-to-drain capacitance *C*_*gd*_ of the middle NMOS transistor which in turn changes the oscillator’s frequency (see refs. [24]–[26] for analytical models of the oscillator’s frequency as a function of circuit parameters, including *C*_*gd*_). Thus, by detecting changes in frequency, we can infer changes in *C*_*coupled*_, which in turn correspond to changes in cell activity.

All 64 sensor pixels share one current reference block where the reference current is established for the selected pixel when the switch XEN closes (Fig. 8). Applying the current as the reference into each pixel significantly reduces the voltage drop caused by the routing metal parasitic resistance. Furthermore, because power is transmitted wirelessly to the chip and the reader antenna imposes a maximum on the power to be dissipated, each pixel is designed to be shut down completely to reduce the power consumption when not selected. This is achieved by adopting a power-gating PMOS between the sensing circuit and the power supply.

The sensor array’s architecture is shown in Fig. 9. The current bias routing *V*_*P BIAS*_ is global because current-driven biases have good immunity against extensive parasitic loading. The selected row address voltage X*<*n*>* goes low to select a specified row while the corresponding gates of the selection transistors in unselected rows remain high. Once a column is similarly selected via Y*<*n*>*, the bias current is passed to the local diode-connected PMOS to supply the selected sensing circuit. As noted above, all unselected sensor pixels are turned off completely. In other words, only when both a specified row and column are selected does the pixel array consume power. Specifically, the sensing oscillator in the selected pixel starts oscillating and sends the output signal to the DOUT pin through the 64-to-1 output YMUX.

**Fig. 9:**
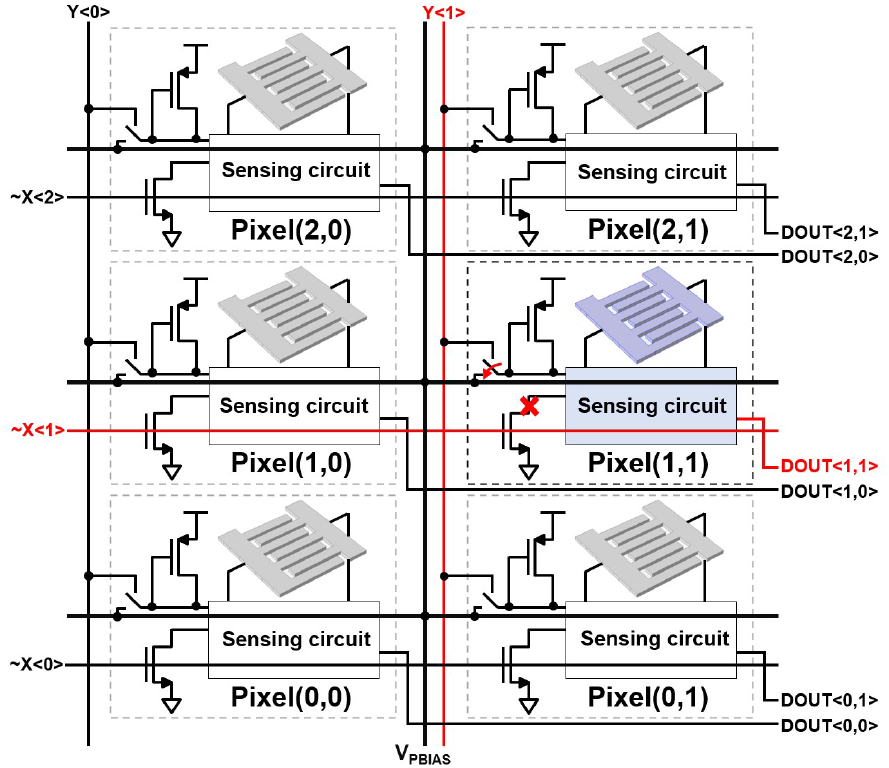
Sensor array architecture. A selected pixel is shown with its selection lines highlighted in red. In this case, the selected pixel address is (1,1), and the global *V*_*P BIAS*_ is used to drive its sensing circuit. The current source has good immunity to the large parasitic loading across the whole array. When a pixel is selected, the sink NMOS is disabled and the current bias is conducted to the sensing circuit.

### C. On-Chip Antenna with Cut Seal Ring

Fig. 10 shows the full symmetric spiral two-loop OCA included in the RFLoC chip. The OCA serves both as an interface for power harvesting and as the antenna for wireless communication. For power transfer, it may be accessed from the chip’s back side, *i*.*e*., through the silicon substrate, or from the front side, *i*.*e*., through the wet test environment. We later show herein that it is preferable to use the back side coupling approach for data transmission.

**Fig. 10:**
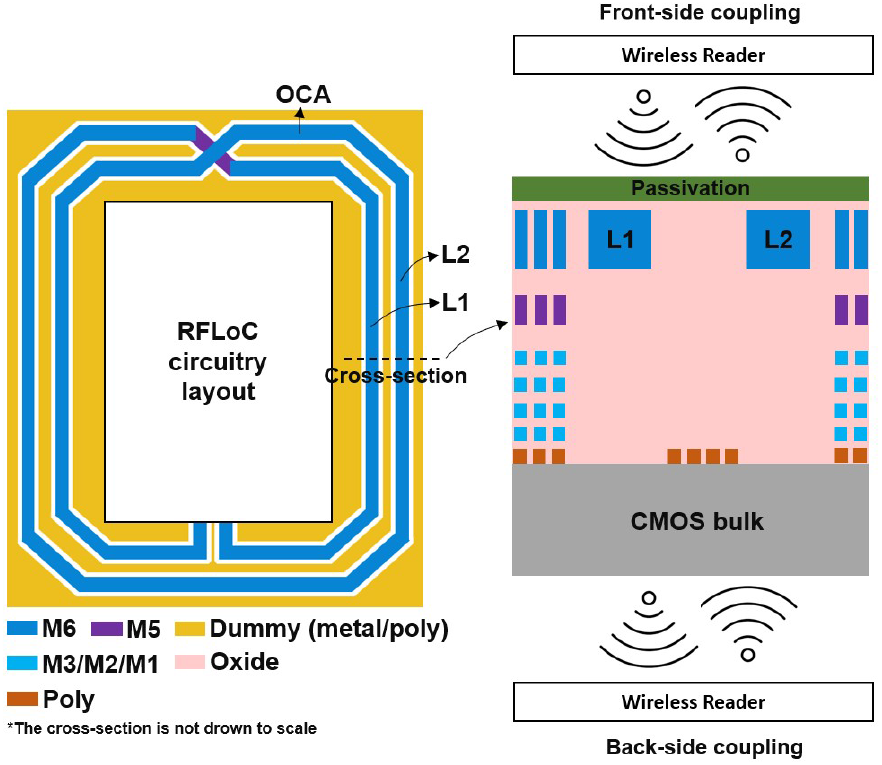
Left panel: schematic of the two-loop OCA used in the RFLoC prototype. The OCA is placed outside of the RFLoC circuitry area and dummy metal layers are excluded under the OCA metal. This enhances the OCA’s self-resonant frequency and reduces interference during back-side coupling. M6 and M5 are thick metal layers that are used reduce the OCA’s resistance. Right panel: Cross-sectional view of the chip under two possible excitation modes (front-side coupling and back-side coupling).

The OCA is implemented using the top thick metal of the process in order reduce the antenna’s AC resistance. Further, to maximize the OCA’s self-resonant frequency and reduce interference from underlying structures, a dummy-metal exclusion layer was used in the layout to ensure that no metal patterns were inserted under the OCA during fabrication. This design choice was limited by the process design rules’ mandated distance between the exclusion layer and the OCA metal. This means that during fabrication, dummy metal patterns could still be placed in the vicinity of the OCA. Nevertheless, as our results show, this did not impede proper operation in either the sense-and-transmit mode or the power harvesting mode.

Equation 2 describes the voltage (*V*_*induced*_) that appears across the OCA and that is induced from the reader wirelessly [27]. Here, *V*_*induced*_ depends on the coupling factor *k*, the OCA’s quality factor *Q*_*OCA*_, the inductance of the reader *L*_*reader*_, the inductance of the OCA *L*_*OCA*_, and on the frequency *f*_*o*_ of the excitation signal. By adopting the ultra-high-frequency band, *i*.*e*., an excitation signal in the 900–920 MHz range, and an OCA implemented in the top thick metal of the process, we were able to achieve a *simulated* quality factor (*Q*_*OCA*_) of 4.65.

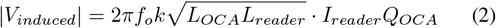

Fig. 11 illustrates a COMSOL™ finite element model that was used to estimate the OCA quality factor that is quoted above. The RF front-end circuitry and the sensor core were modeled as metal structures that were placed within the symmetric two-loop OCA. Signal and power connection wires were routed under the OCA, but these wires were shielded with grounded metal planes disposed between the OCA and the wires. The chip’s seal ring was cut, in order to avoid any parasitic loading to the OCA, as explained below. Furthermore, given the thickness of the metal, which was obtained from foundry-provided data and the dimensions of the OCA, its inductance was estimated by the model to be 17.53 nH, and its quality factor was calculated to be 4.65.

**Fig. 11:**
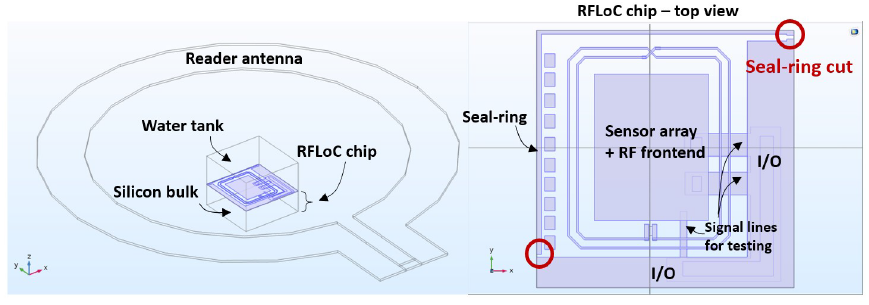
Overview of the finite element model used to evaluate the OCA’s quality factor and inductance. The chip’s main constitutive circuits are modeled as a single metal plate with connecting wires running under the OCA. The chip’s seal ring is modeled using a discontinuous metal loop (indicated by the seal ring cut).

The OCA’s coupling factor *k* was modeled as shown in Equation 3 where *x* is the distance between the OCA and the reader antenna, and *r*_*OCA*_ and *r*_*Reader*_ are the radii of the OCA and the reader antenna, respectively. As indicated by Equation 3, *k* depends strongly on both antennas’ geometries and on the coupling distance [28]. For instance, the coupling becomes stronger with a shorter distance between the two antennas. We note that under front-side coupling excitation, *x* is limited by the culture well’s height. This restriction means that the applicable antenna aperture is also limited. For example, for a 5 to 10 mm height well, *r*_*Reader*_ should be close to that height value in order to achieve a higher coupling factor, which in turn results in a higher induced voltage on the chip.

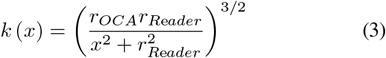

Our finite element model revealed that the chip’s seal ring significantly reduced the coupling factor *k* because it formed a larger parasitic loop placed outside of the OCA. This means that when the chip is excited by the reader antenna’s RF signal, the seal ring deteriorates the wireless power transfer efficiency as a result of the RF energy being coupled to this parasitic loop.

Seal rings are added to prevent stresses from reaching sensitive circuits during the CMOS fabrication process. They are usually implemented using a fully connected metal stack forming a loop at the chip’s perimeter. Seal rings are mandatory in multi-project wafer runs and thus cannot be removed by the designer, as would be ideal for our application. As an aside, we note that the seal ring also poses significant challenges to integrative packaging methods that rely on planar metal redistribution. Specifically, the seal ring can cause discontinuities in the metal redistribution as a result of the ring’s surface topography [29].

We mitigated the seal ring’s parasitic effects by “cutting” it. This was done with the foundry’s agreement, and it was achieved simply by making the seal ring discontinuous using a small cut in the metal pattern layout. Our model revealed that for a 10 mm diameter reader antenna at 1 mm distance, the coupling factor *k* improved by 52%, *i*.*e*. from 0.015 to 0.0228, by breaching the seal ring’s continuity with a cut in the metal stack layout.

### D. Wireless Power Management

Fig. 12 illustrates the rectifier topology that was used to convert the induced RF voltage in the OCA (*V*_*induced*_) to a DC voltage (*V*_*DD*_) that supplies the sensor system and the various other on-chip circuitries. As previously noted, the amplitude of the inductively coupled RF input voltage is a strong function of the distance between the external reader antenna and the OCA. In order to allow a wider operation range, a CMOS rectifier with a gate boosting scheme [30] was adopted in order to reduce the start-up voltage and improve efficiency under low RF input voltages that would result from the two antennas being far apart.

**Fig. 12:**
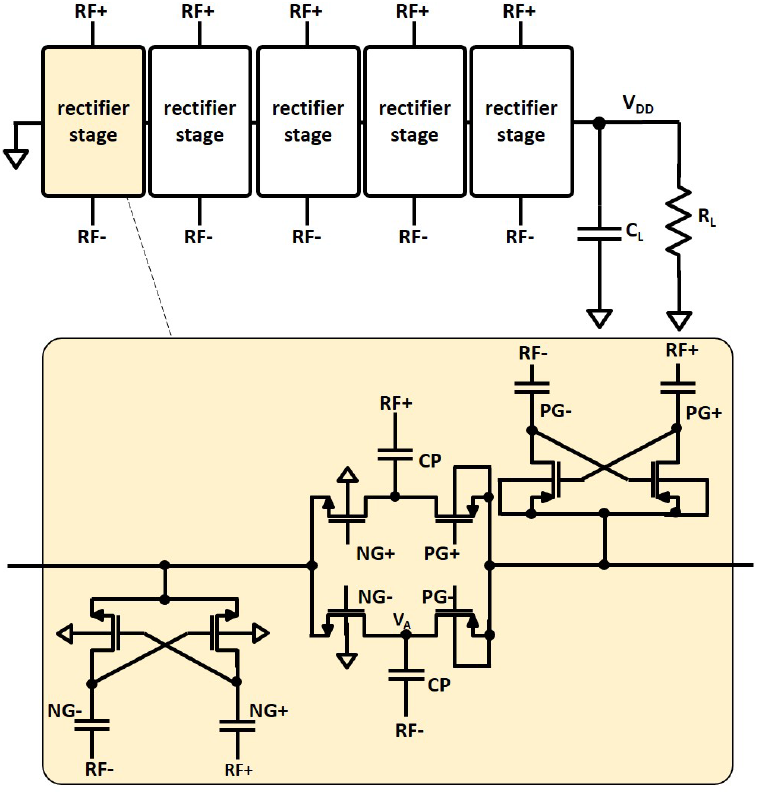
Five-stage boosting rectifier circuit topology. This rectifier design increases the conversion efficiency for lower RF input amplitude voltages.

The boost circuit does not need extra biases to be driven compared to conventional rectifier topologies [31]; instead, it is charged by the input and output node voltages of each stage. The gate-boosting pairs couple both PMOS (*PG*+, *PG−*) and NMOS (*NG*+, *NG−*) gate voltages to lower and higher levels, respectively, because the outputs of the boost pairs only see the gate and junction capacitances instead of the large loading current. This feature reduces the on-resistance of the MOS devices so that charges can be drained to the output node more efficiently. The simulation results show that *V*_*DD*_ can be boosted by 200 mV for a 600 mV RF input voltage.

The output of the rectifier is connected to a shunt regulator in order to control the DC voltage level. The shunt regulator topology utilized is shown in Fig. 13. When *V*_*DD*_ exceeds the desired level, *V*_*D*_ will be higher than *V*_*REF*_ so that the discharge path activates to stabilize *V*_*DD*_. Nevertheless, considering the gain of the amplifier’s error, additional stability features are needed for the regulator in order to ensure the chip’s proper operation in different modes.

**Fig. 13:**
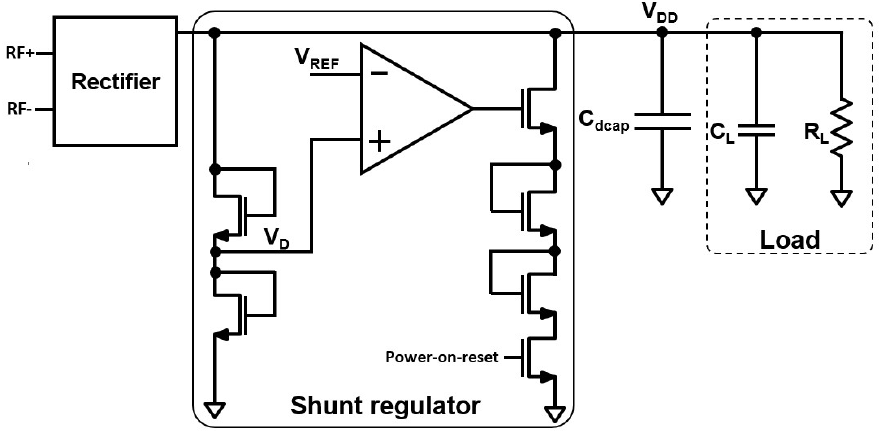
Shunt regulator topology. The shunt regulator is enabled after power on. *V*_*D*_ samples *V*_*DD*_ using a MOS voltage divider. The regulator compares *V*_*D*_ and *V*_*REF*_ to set the regulated *V*_*DD*_ level.

For example, the loading current of the sensor-enable period is higher than that of the standby period, yielding different stability constraints. As such, to prevent abnormal current sinking when *V*_*REF*_ has not established a stable value during wireless power-up, the sinking path is gated by a power-on-reset signal. Further, to prevent stressing the transistors comprised in the sensor core and the RF-front end, the target regulated voltage is set at 2 V. Lastly, two diode-connected NMOS devices are placed along the sinking path to prevent instantaneous *V*_*DD*_ drops during power-on.

Another important feature for power management is the bias circuit that is used to provide a reference current that is distributed to a pixel during a sensing operation. This circuit is shown in Fig. 14. It is a constant gm-bias reference, and it is equipped with a continuous-time startup circuit.

**Fig. 14:**
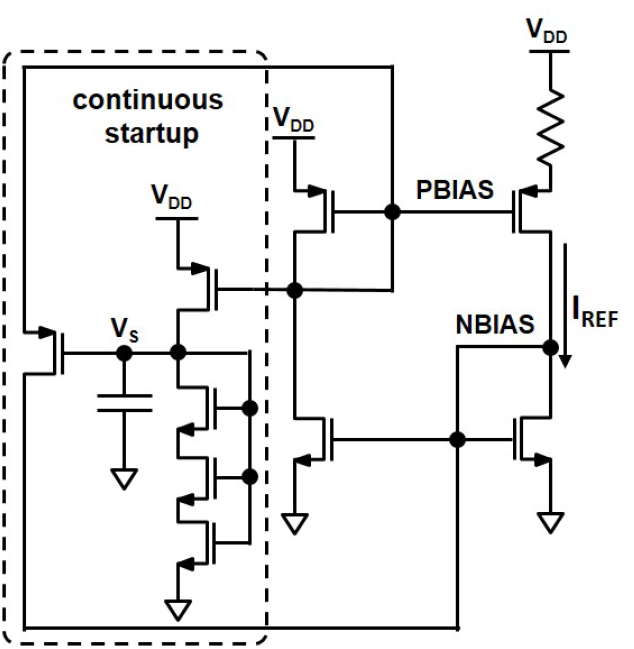
A constant gm-bias circuit with a continuous time start-up circuit is used to protect against *V*_*DD*_ brownouts.

The gm-bias reference current *I*_*REF*_ must remain stable in the presence of *V*_*DD*_ fluctuations. One cause of fluctuation may be a variation in the distance between the two antennas as a result of a user disturbing the two boards. A fluctuation can happen because the *V*_*DD*_ level depends on the distance between the two antennas, the angle of incidence of the RF energy, as well as the transmitted signal power from the external antenna. Therefore, during operation, a variation in any of these parameters can weaken the coupling between the reader antenna and the OCA, resulting in an appreciable change in *V*_*DD*_.

These conditions can cause the internal voltages of the gm-bias circuit to drop, which would shut down all the devices, causing an interruption in the generation of *I*_*REF*_ . Even when *V*_*DD*_ is quickly recovered, the constant gm-bias circuit may keep turning off if the start-up circuit can not be triggered again. Thus, to prevent the constant gm-bias reference circuit from locking after a short brownout, the continuous-time startup circuit continually sinks a small current to detect the status of the gm-bias circuit’s internal biases such that there is no dead-zone for the startup circuit. With the continuous-time start-up circuit, when a brownout in *V*_*DD*_ happens, *I*_*REF*_ may drop to zero, but the internal voltages may still be at non-zero levels. If *V*_*DD*_ is recovered after the brownout, the gm-bias is able to start up without locking into the shut down region, independently of the nodal voltages.

### E. Data Uplink Transmission

In the sense-and-transmit mode, the sensor core’s output data are wirelessly transmitted to the external reader antenna via a data uplink module. Our data uplink design adapts the on-off keying (OOK) modulation scheme [32]. The OCA can operate normally during the OOK scheme’s on state. In the off state, the *RF* + and *RF −* ends are shorted. This means that when the OCA switches from the on state to the off state, its resonance frequency is shifted from the carrier frequency, and the back-scattered data can be sensed by the reader antenna [27], [33].

However, the off state also terminates wireless power transfer to the chip from the reader. This situation risks creating a significant drop in *V*_*DD*_ that could in turn shut down the sensor system and other critical circuits. Hence, in our implementation of the OOK scheme, the period of the off state is controlled precisely, and a large decoupling capacitor is used to stabilize the *V*_*DD*_ power supply against the transients that are generated as a result of the uplink transmitting data. To further ensure proper operation, each logic 1 one bit outputted from the PISO interface triggers one transmission pulse TX from the pulse generator (Fig. 15). The transmission pulse width is controlled so as to be long enough to be distinguishable in the received signal at the reader but short enough to maintain the *V*_*DD*_ level.

**Fig. 15:**
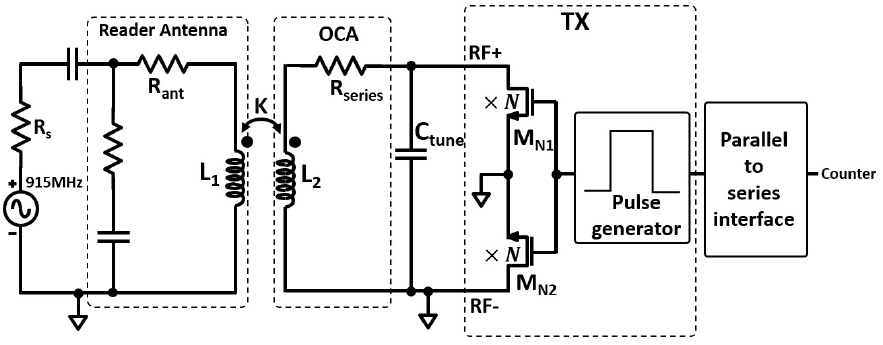
Transmitter (TX) architecture. The transmitter can alter the resonance frequency of the wireless link, resulting in reduced coupling, which in turns can cause *V*_*DD*_ to drop. As such, the transmission pulse width is controlled precisely by the pulse generator to minimize potential adverse effects on *V*_*DD*_ when transmitting data.

### F. Sensor Data Modulation

The output signal from a selected sensor pixel is an oscillating signal, and it is fed to a ripple counter (Fig. 5). This is done to count the number clock edges that occur during the enable period of the output counter, which defines the measurement time *T*_*meas*_. The number of clock counts registered during the period is correlated with a change capacitance at the interface. When the output counter is disabled, the number of registered clock counts is stored in the output counter’s registers automatically, and it is ready to be used for the next stage. Since the current bias controls the sensor’s output frequency, that frequency is designed to be within the operational range of the output ripple counter to prevent counting errors.

The output counter is a 24-bit ripple counter. However, the OOK modulation transmitter can only transmit one-bit per cycle as described previously; therefore, a PISO interface is employed after the counter to convert the 24-bit parallel output data into a series output data stream. When the asynchronous PISO enable signal is asserted, a fetch signal synchronizer circuit (Fig. 16) generates a pulse signal (FETCH) that lasts two clock cycles to ensure that the PISO’s shift registers can fetch the counter’s data.

**Fig. 16:**
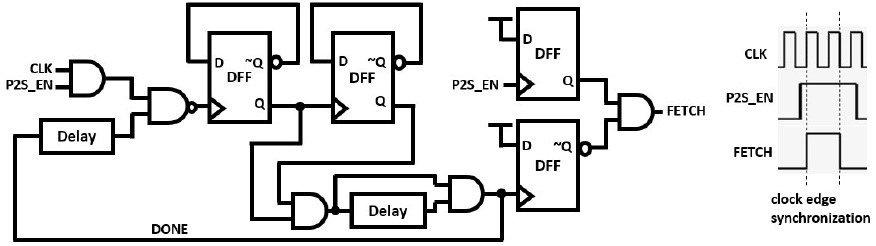
The fetch signal synchronizer circuit is configured to copy the counter data into the PISO shift registers. The synchronizer guarantees that the FETCH signal keeps high across at least one clock edge.

Fig. 17 shows the connections between the output ripple counter and the PISO’s shift register. To demarcate the first and last bits of the 24-bit series output data stream, dummy patterns were inserted before and after the serial output sensor data. This was achieved by adding extra registers with hardwired data representative of the dummy data at the PISO’s preamble stages.

**Fig. 17:**
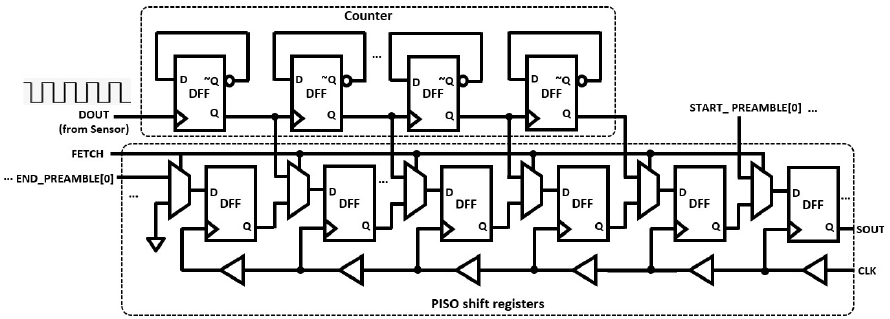
The PISO shift registers contain start/end dummy data that are inserted at the beginning and end of the serial counter data.

### G. Chip-on-PCB Mounting and Device Packaging

Fig. 18 shows a photomicrograph of the manufactured RFLoC CMOS chip bonded on a PCB. The chip’s I/O bond pads are connected to the PCB’s soft gold-finished pads using aluminum bonding wires. A water-impermeable epoxy (Durapot 865, Cotronics Inc.) was used to coat all the bonding wires in order to protect the aluminum from the liquid analyte. The epoxy was applied such that the sensor array and the OCA remain exposed.

**Fig. 18:**
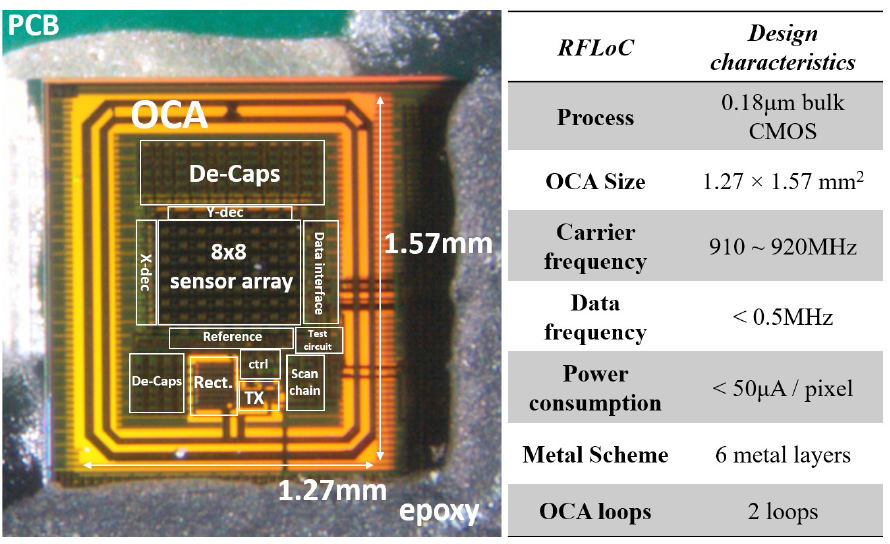
Photomicrograph of the RFLoC sensor chip packaged and partially encapsulated on a printed circuit board. A summary of the chip’s characteristics are shown to the right of the photograph.

The RFLoC chip was manufactured in a six-metal 0.18 *µm* commercial CMOS process with two top thick-metal layers. The thick-metal layers were used to construct the OCA to reduce its resistance at high frequencies. The current dissipated in each pixel during operation was less than 50 *µA*. The dimensions of the RFLoC chip (without the I/O pads’ area) were 1.27 *mm ×* 1.57 *mm*. Each pixel was 30 *µm ×* 30 *µm* in size, and the 8 × 8 array in the middle of the chip was 470 *µm ×* 606 *µm* in size.

## III. Experimental Constructs

### A. Antenna Coupling Methods

In a use case, the RFLoC chip will be interfaced with a wet test environment provided by a culture well filled with a liquid medium. The chip will further be interfaced with external equipment for data collection and processing and with a microscope for ground-truth imaging data used to correlate the measured capacitance data with biological activity.

To that end, two methods may be used to couple the reader antenna to the OCA. The first method consists of placing the reader antenna on the front side of the chip, *i.e*., above the culture well. The second method consists of placing the reader antenna under the PCB to couple to the OCA from the back side of the chip.

The front side setup is shown in Fig. 19. In this approach, the culture well’s height is the limiting factor as it readily imparts a minimum allowable distance for positioning the reader antenna. Furthermore, in order to allow microscopy to take place, the reader antenna must be hollowed in order to create an optical path for the microscope objective; this requirement introduces additional complexity to the prototyping process.

**Fig. 19:**
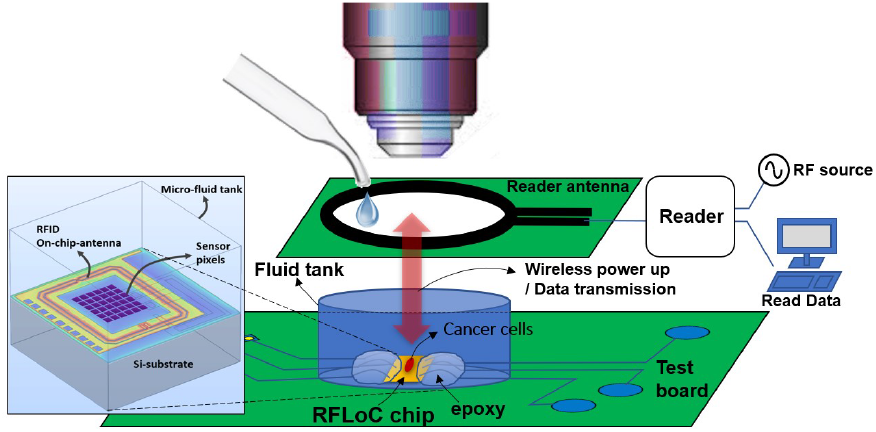
Front side excitation and uplink. The reader antenna is placed above the chip, and it forms the link at the chip’s front side, through the culture medium.

Nevertheless, despite these drawbacks, the front side setup is advantageous in situations where devices cannot be placed under the chip. For example, such a situation may arise when the chip is placed on the bottom surface of a cell culture incubator and space is limited under the board to insert the reader antenna.

The back side setup is shown in Fig. 20. By putting the antenna under the RFLoC chip and the PCB substrate, the chip may be used in a conventional manner when there are no restrictions for the placement of the reader antenna.

**Fig. 20:**
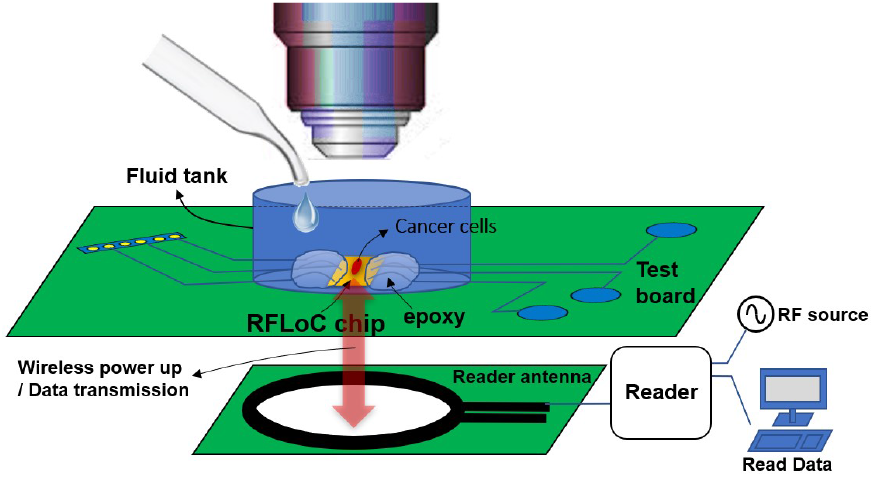
Back side excitation and uplink. The reader antenna is placed below the chip, and it forms the link at the chip’s back side, through the PCB and silicon substrate.

### B. Wireless Reader Design

The wireless RFID reader’s block diagram is shown in Fig. 21. The reader is configured to transmit the UHF RF signal and to receive the uplink data from the RFLoC chip. By design, the nominal frequency of the RF carrier signal is between 900 to 920 MHz, *i.e*., within the ISM band. An RF signal generator is used to generate the UHF RF signal. Using an RF power splitter, the generated signal is passed to both a receiving mixer and to a power amplifier having a gain of 25 dB.

**Fig. 21:**
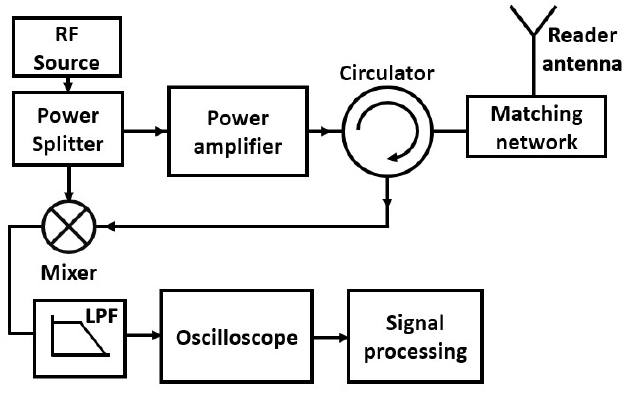
Wireless reader block diagram. The RF carrier signal is between 900 and 920 MHz, *i.e*., within the ISM band.

An RF circulator is applied between the reader antenna and the power amplifier to send the RF signal into the reader antenna while also maintaining a path for receiving the uplink data. The selected circulator has high isolation (25 dB to 30 dB) between its two adjacent ports. The received uplink signal is fed into the mixer to distinguish the signal from the carrier. Subsequently, the mixer’s output signal is fed to a low pass filter and sent to an oscilloscope for viewing and analysis. Additional signal processing was performed in MATLAB™ in order to retrieve the uplink digital data. Figs. 22 and 23 shows our test bench, using each of the two coupling methods discussed above. A National Instruments data acquisition system was used to control the chip and test its internal signals via the I/O pads for design verification purposes.

**Fig. 22:**
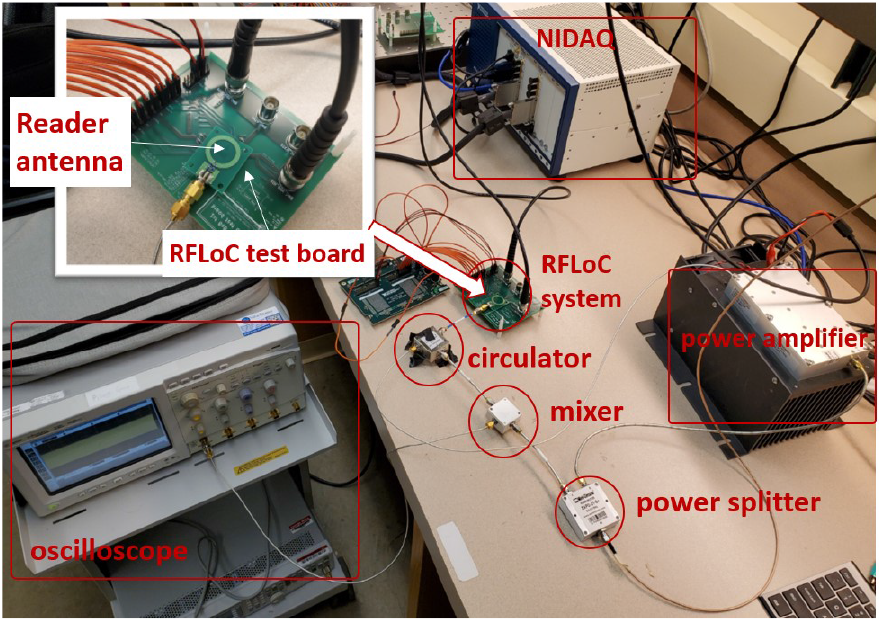
Test setup for the RFLoC test board and the wireless UFH reader. The front side setup is applied in this demonstration without the microscope.

**Fig. 23:**
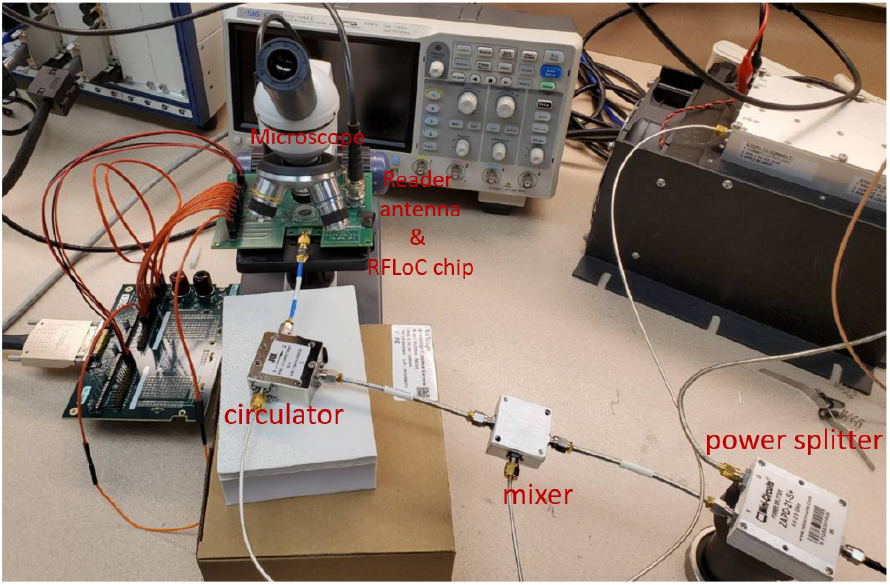
Test setup for the back side excitation and uplink configuration. Here, the placement of the reader antenna does not interfere with the microscope’s optical path.

**Fig. 24:**
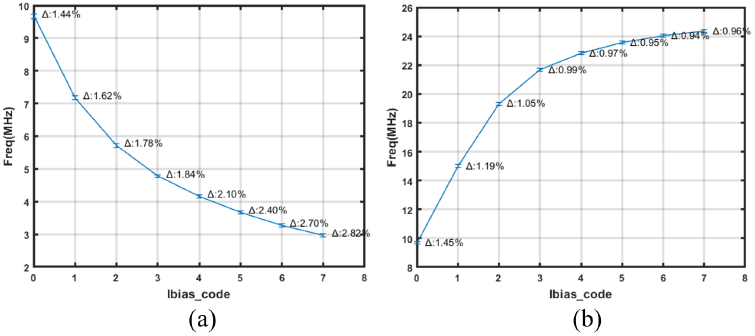
Pixel base frequency tuning by programming the PMOS current mirror. The frequency decreases with increasing code value.

**Fig. 25:**
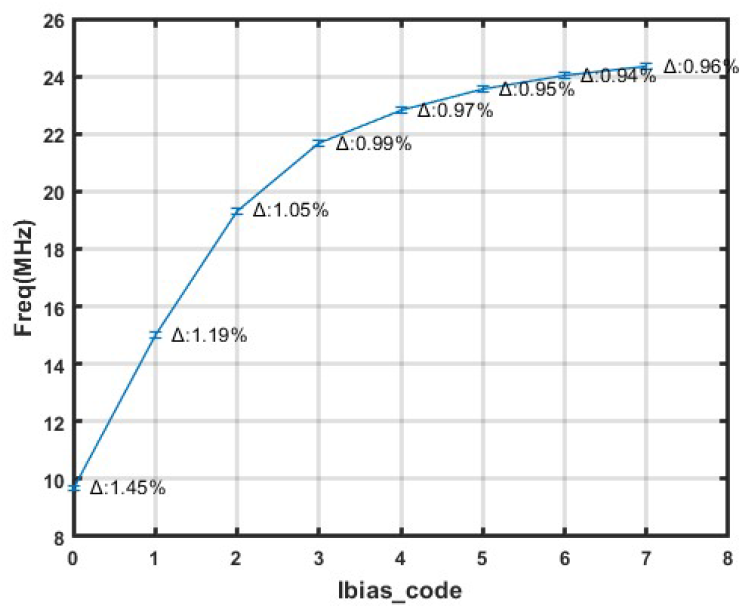
Pixel base frequency tuning by programming the NMOS current mirror. The frequency increases with increasing code value.

## IV. Measurement Results

In this section we report characterization results of the various RFLoC functions described above. First, we report on the sensor core’s performance featuring results from pixel frequency tuning and analyte characterization experiments. We then report on the chip’s wireless power transfer efficiency.

Lastly, we report characterization results of the chip’s uplink module under each of the front and back side coupling modes mentioned above.

### A. Pixel Frequency Tuning

The frequency of each pixel in the 8×8 capacitance sensor array can be tuned by changing the control codes in the bias generator (Fig. 8). The tuning codes can be programmed to increase or decrease the oscillator’s base frequency by switching the PMOS current-mirror options or NMOS current-mirror options, respectively. Figs. **??** and 25 show the measured results of tuning the oscillator’s frequency. In sum, the base frequency can vary from *∼* 3 MHz to *∼* 24 MHz.

The measured maximum percentage change in frequency for all of the 64 pixels and among all codes was *∼*2.8%, which corresponded to a maximum standard deviation of 10^*−*3^ MHz. Figs. 26 and 27 show the standard deviation of the frequency across the array as a function of the selected bias code, whether the frequency increases with the control codes or whether it decreases. To further provide flexibility in tuning the pixel’s base frequency, the RFLoC prototype was equipped with an I/O pad onto which a specific bias current could be forced externally. The data from Figs. 26 and 27 suggest that a specific programmed bias current results in a unique standard deviation across the array. As such, we also expect the same relationship for currents that are forced through the I/O pad.

**Fig. 26:**
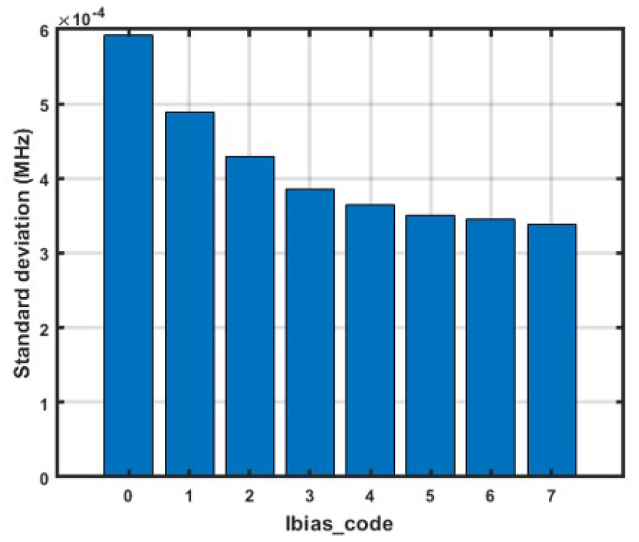
Standard deviation of the frequency across the 8 × 8 sensor array when tuning is achieved using the PMOS current mirror option to decrease the pixel frequency with increasing values of the bias code.

**Fig. 27:**
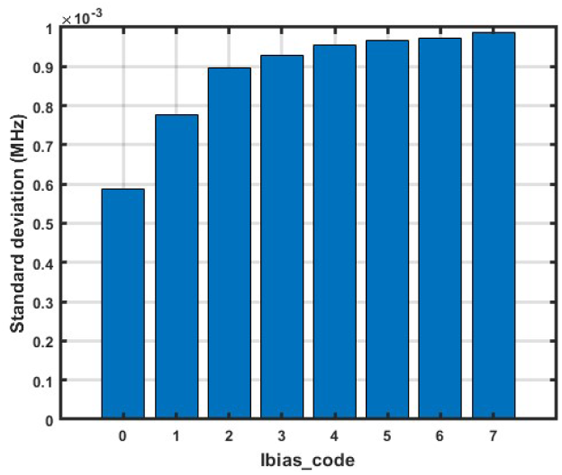
Standard deviation of the frequency across the 8 × 8 sensor array when tuning is achieved using the NMOS current mirror option to increase the pixel frequency with increasing values of the bias code.

### B. Analyte Characterization

To validate the sensor core’s function, three liquid analytes were used. They were: ethyl alcohol (62%), water, and coffee. The change in frequency resulting from these analytes was referenced to air, *i.e*., to the frequency registered with no analyte present on the chip.

As shown in Fig. 28, the average frequencies measured across the array for air, water, ethyl alcohol, and coffee were 9.68, 9.27, 9.13 and 9.06 MHz, respectively. Fig. 29 shows the bitmaps representing the spatial distribution of the frequencies across the array at different bias conditions for measurements of air and water. Fig. 30 shows the same data when the array is partially covered with water.

**Fig. 28:**
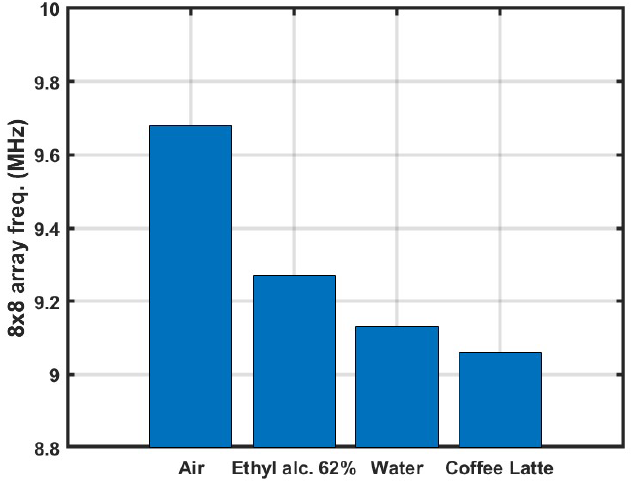
Average frequency of the 8× 8 sensor array for different analytes measured in reference to air and at room temperature.

**Fig. 29:**
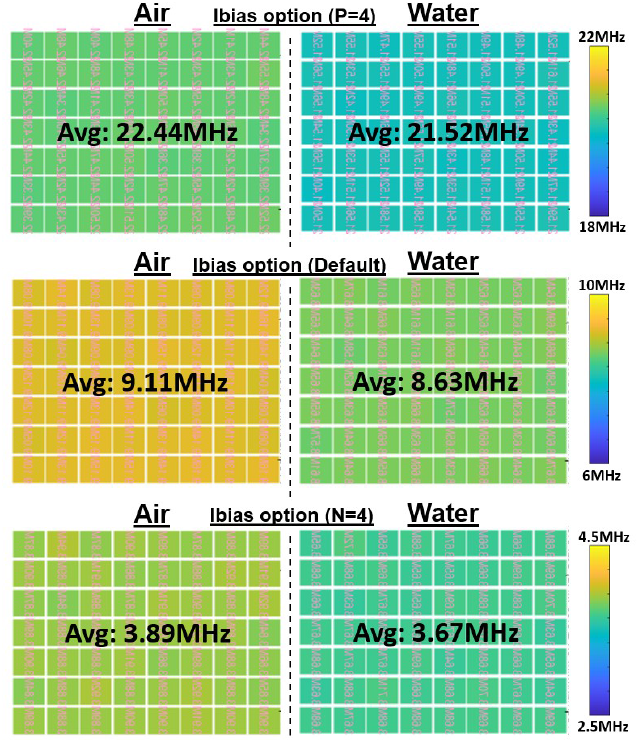
Bitmaps for water and air measurements using different bias current options.

**Fig. 30:**
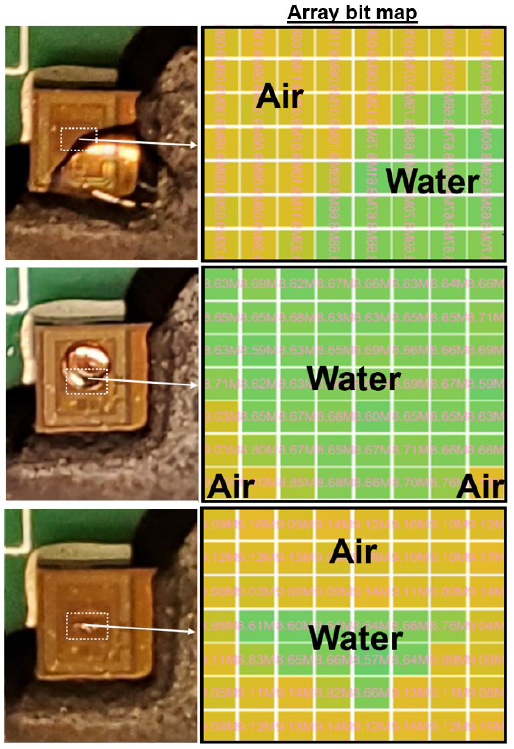
Photomicrograph and capacitance images of water partially covering the RFLoC sensor array. The measurements were made for a fixed bias current options. The bitmaps show that the pixels covered by the drop of water have lower frequencies than the pixels exposed to air. The spatial boundary from the capacitance image indicates the drop’s perimeter.

### C.*Wireless Power Transfer*

Fig. 31 shows the front and back-side setups that were used to characterized the chip’s performance in the harvesting mode. The RF carrier’s amplitude generated by the source signal generator determined the transmitted electromagnetic wave’s strength. The rectified *V*_*DD*_ in the harvesting (standby) mode and in the sense-and-transmit (pixel-on) mode were measured for cases where the RFLoC chip was placed in air or water was used as the analyte. These data are shown in Figs. 32 and 33.

**Fig. 31:**
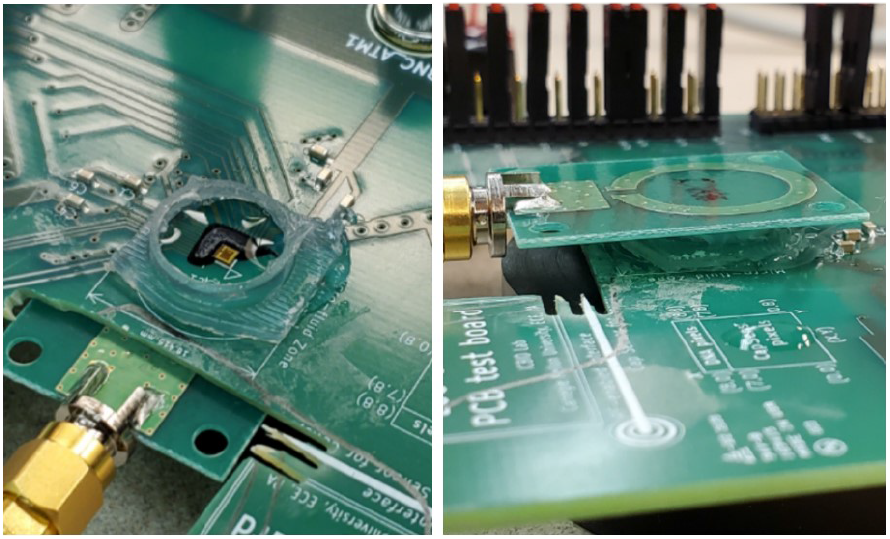
Experimental setups for characterizing the wireless power transfer efficiency using the back-side-setup (left) and front-side-setup (right). A 4 mm-height plastic well encloses the RFLoC chip and serves to contain a liquid analyte (*e.g*., water).

**Fig. 32:**
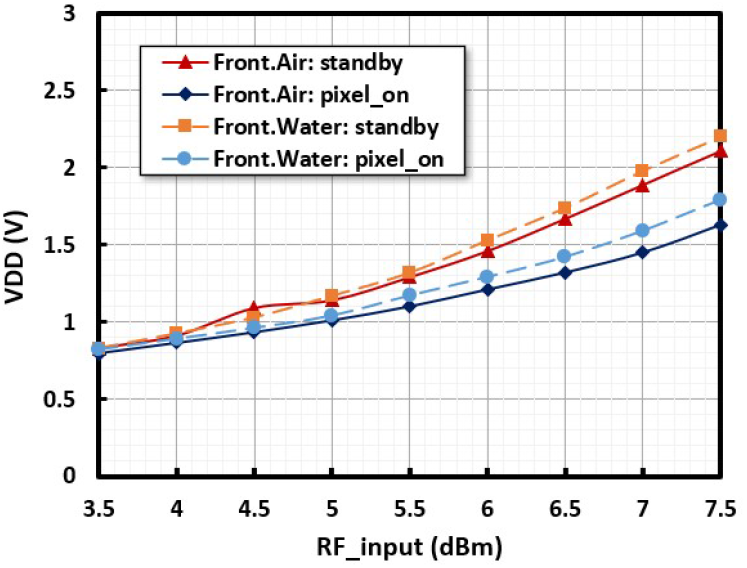
*V*_*DD*_ as a function of the reader’s input RF amplitude at 4 mm distance between the reader antenna and the RFLoC chip’s OCA. The measurements were made in the front side coupling mode. *V*_*DD*_ drops when the pixel array is activated, *i.e*., when the chip is switched from the harvesting (standby) mode to the sensing mode.

**Fig. 33:**
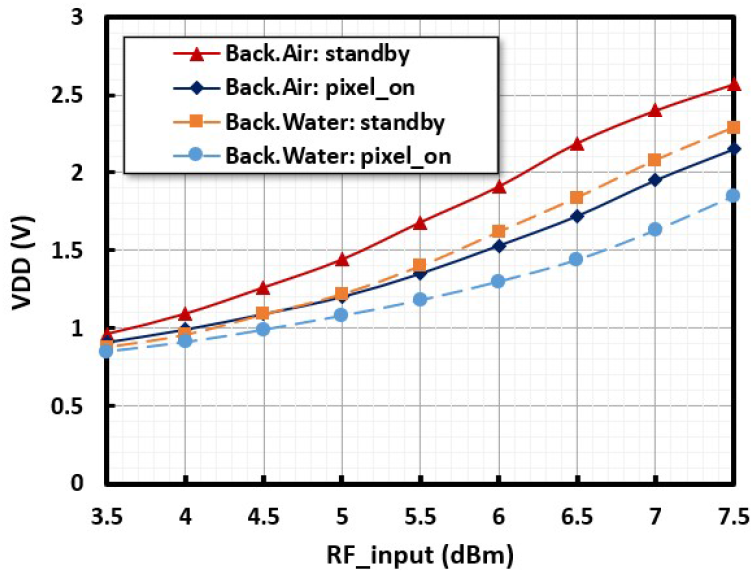
*V*_*DD*_ as a function of the reader’s input RF amplitude at 4 mm distance between the reader antenna and the RFLoC chip’s OCA. The measurements were made in the back side coupling mode. *V*_*DD*_ drops when the pixel array is activated, *i.e*., when the chip is switched from the harvesting (standby) mode to the sensing mode.

In the standby mode, the bias voltage, bias current, and *V*_*DD*_ were fixed and stable, and the pixel array was shut down, whereas in the pixel-on mode, the pixel array was continuously enabled during measurement. The higher the RF amplitude the higher the *V*_*DD*_ induced on the chip. Furthermore, as expected, with both water and air test cases, *V*_*DD*_ dropped when the pixel array was activated because the operating sensor array consumed power. The distance between the reader and the RFLoC chip was 4 mm, and it was limited by the height of the well on the front side. In the back side coupling mode, the limitation was imparted by the PCB’s thickness. Nevertheless, for consistency, the reader antenna was placed at a distance of 4 mm from the PCB’s bottom surface.

With respect to the induced *V*_*DD*_, we found that the wireless power transfer performance for the front side coupling mode (Fig. 32) was similar when the chip had no analyte on its surface (*i.e*., in air) to when water was placed in the culture well. We surmise that the ability of inducing the adequate *V*_*DD*_ in the front side coupling mode in the presence of the liquid medium was not compromised because of the proximity of the two antennas. Nevertheless, we note that the overall *V*_*DD*_ of the back side coupling mode is higher than the front-side-setup, and thus, the back side setup is preferable. Fig. 33 shows *V*_*DD*_ as a function of the RF amplitude when the reader antenna is placed on the back side.

Equation 4 shows how we calculated the RFLoC chip’s wireless power transfer efficiency (PTE). The load resistance to the OCA was estimated to be 10 *k*Ω, and we used a maximum *V*_*DD*_ of 1.3 V and an RF carrier strength of 6 dBm for the calculation. Further, the power amplifier had a gain of 25 dBm, and the total loss of the reader components was estimated to be 7.2 dB. These figures resulted in a radiated power of 23.8 dBm (0.24 W), yielding a PTE of 0.07%. The gain and loss budget for the reader antenna are shown in Fig. 34.

**Fig. 34:**
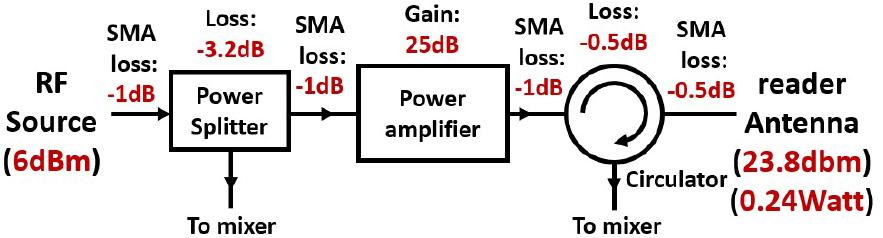
Gain and losses in the signal path of the reader antenna. The radiated power was estimated to be 23.8 dBm (0.24 W).

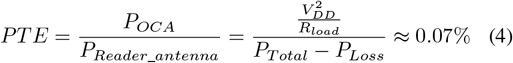

### D. Uplink Measurements

Fig. 35 shows the transmitted uplink data that the reader receives from the RFLoC chip. The chip senses and records data from the selected pixel output for a given sensor enabling *T*_*meas*_ time, and it transfers the data out through the transmitter as previously shown in Fig. 15. To distinguish the received signal from the carrier signal and to indicate the location of the sensor data, the 24-bit output result is transmitted with a bit dummy data preamble (11100101)_2_ and an 8-bit dummy data appendix (11000101)_2_. The data shown in Fig. 35 are for the front side coupling mode, using air as a test case, and the sensor output data is (11111011000)_2_, which corresponds to m2008 counts out of 2^24^ *−* 1 maximum possible counts given the output counter’s bit depth. The top panel shows the original data stream that was received, stored at the oscilloscope, and retrieved using a PC. The carrier frequency was 915 MHz. The uplink signal was demodulated from the oscilloscope data using a MATLAB™ signal processing routine. The bottom panel shows the received data stream after demodulation and signal processing.

**Fig. 35:**
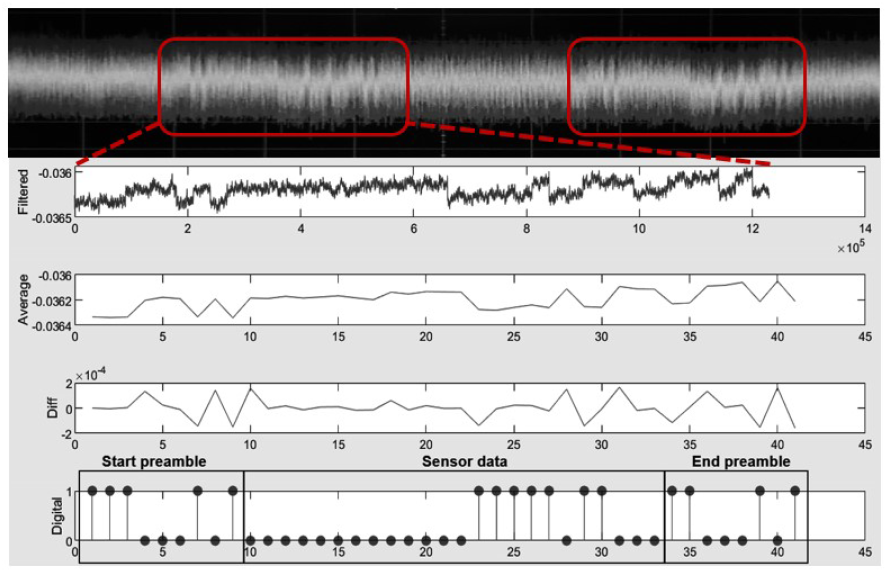
Wireless uplink data received from the RFLoC chip. The red rectangles identify data markers which surround the sensor output data. The data stream includes the 9-bit preamble pattern, the 24-bit sensor output data, and the 8-bit appendix pattern.

Our measurements revealed that wider transmission pulse widths (TX) achieved higher signal-to-noise ratios. For instance, in our uplink characterization experiments, we used TX pulse widths of 125 ns, 250 ns, 500 ns, and 1 *µ*s. The data show that the pulse width may significantly affect transmission.

For example, in Fig. 36a, which show measured data from utilizing the front side coupling for air, we fed a pattern of synthetic data to the uplink module via a dedicated I/O pad. Considering the top chart, which corresponds to a TX pulse width of 125 ns, the test pattern was not readily identifiable. As the TX pulse increased, the received signal began to show the full input sequence. The pattern was clearly visible for a TX pulse width of 1 *µ*s. In such a case, no additional signal processing beyond demodulation would be necessary to retrieve sensor data transmitted from the uplink, contrary to the case depicted Fig. 35, which used a TX pulse width of 100 ns to transmit the sensor data to the reader.

**Fig. 36:**
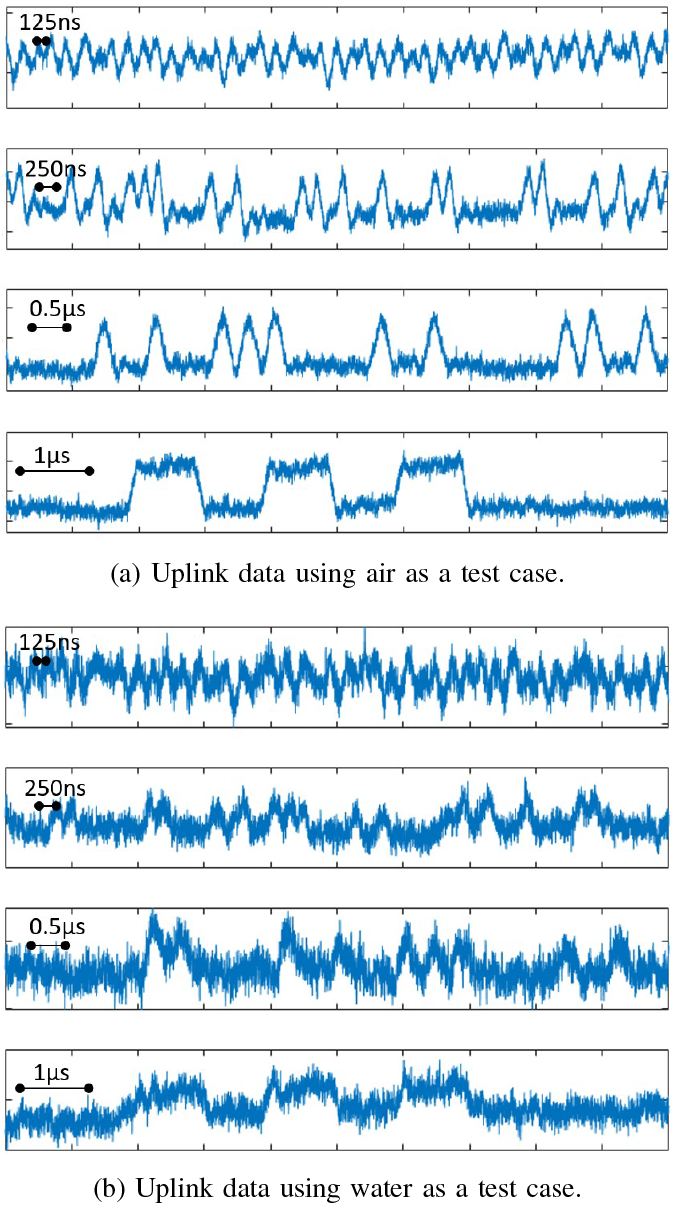
Uplink measurement waveforms using the front side coupling mode. The TX pulse width is set to 125ns, 250ns, 0.5*µ*s, and 1*µ*s.

The measured data further revealed that in the front side coupling method, additional signal processing could still be needed when there is an analyte present, even for wide pulse widths. This is shown in Fig. 36b where the same trend with pulse width is observed, but in contrast, for a pulse width of 1 *µ*s, there is significantly more noise compared to the case where there is no analyte on the chip. We note that no significant differences were observed in the harvesting mode when an analyte was present on the chip. As such, the present results are thus contrary. In other words, in the transmission mode, there is a clear degradation of SNR when an analyte is present on the chip.

We posit that this scenario is the result of conduction paths in the water (likely resulting from ionic species) that induce substrate eddy currents which effectively lowers the coupling factor between the two antennas [34]–[36]. Nevertheless, in our application, the uplink signal from the front-side setup is still observable if the the TX is pulse width is long enough.

We caution that with cell media, where there can be a more significant ionic component than the water analyte used here, we do expect the ability to induce the proper *V*_*DD*_ and the ability to transmit data via the uplink module to be reduced in the front-side coupling mode. Any reduction in performance may be compensated by increasing the RF carrier’s amplitude and by minimizing the distance between the two antennas.

The back side coupling mode addresses the limitations of the front side coupling mode. For example, as shown in Fig. 37a and Fig. 37b, the same conclusion holds with respect to the uplink’s SNR as a function of the TX pulse width. Specifically, longer pulse widths obviate the need for additional signal processing. However, in the case of a liquid medium being present on the chip, the signal is much more readily observable at long pulse widths. This is because the liquid medium does not obstruct the wireless link between the OCA and the reader antenna [37]. Therefore, the back side setup’s SNR is better than that of the front side setup.

**Fig. 37:**
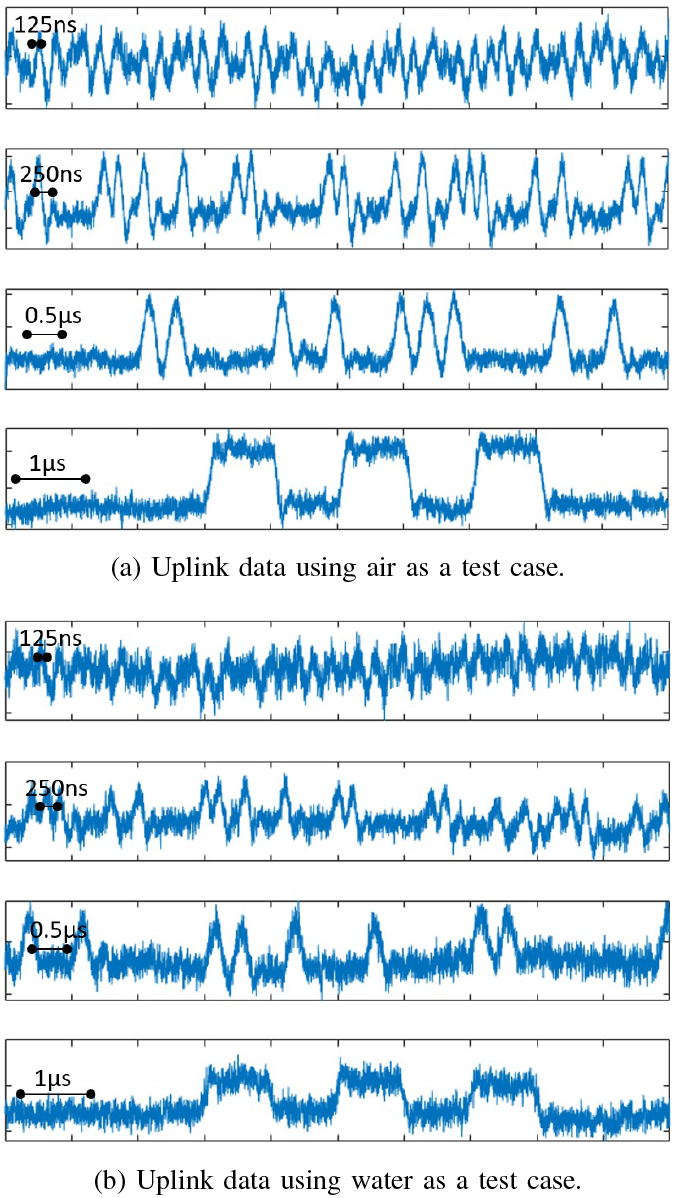
Uplink measurement waveforms using the back side coupling mode. The TX pulse width is set to 125ns, 250ns, 0.5*µ*s, and 1*µ*s.

Generally, in the back side coupling mode, the distance between the two antennas is limited only by the thickness of the PCB and by the RFLoC chip’s thickness. In our case, the PCB’s thickness was 2.5 mm, but it possible to obtain PCBs that are as thin as 0.2 mm. Furthermore, the chip’s thickness is normally less than 0.5 mm. Therefore, in the back side coupling mode, the distance can be precisely controlled to be within 3 mm. Finally, since in either one of the cases discussed above the *V*_*DD*_ supply still holds when the TX pulse width is made to be as long as 1 *µ*s, we set the nominal pulse width for transmission at that value in order to receive a stable uplink signal.

## V. Conclusion

We compared several capacitance sensing platforms with the proposed RFLoC chip. Performance metrics and design features for these devices are reported in Table I along with those of the prototype featured herein. Three distinct architectures were compared. They were the charge sharing (CS) architecture, the charge-based capacitance measurement (CBCM) architecture, and the capacitance to frequency (CF) architecture, the latter being the configuration of the RFLoC chip.

**Table 1.**
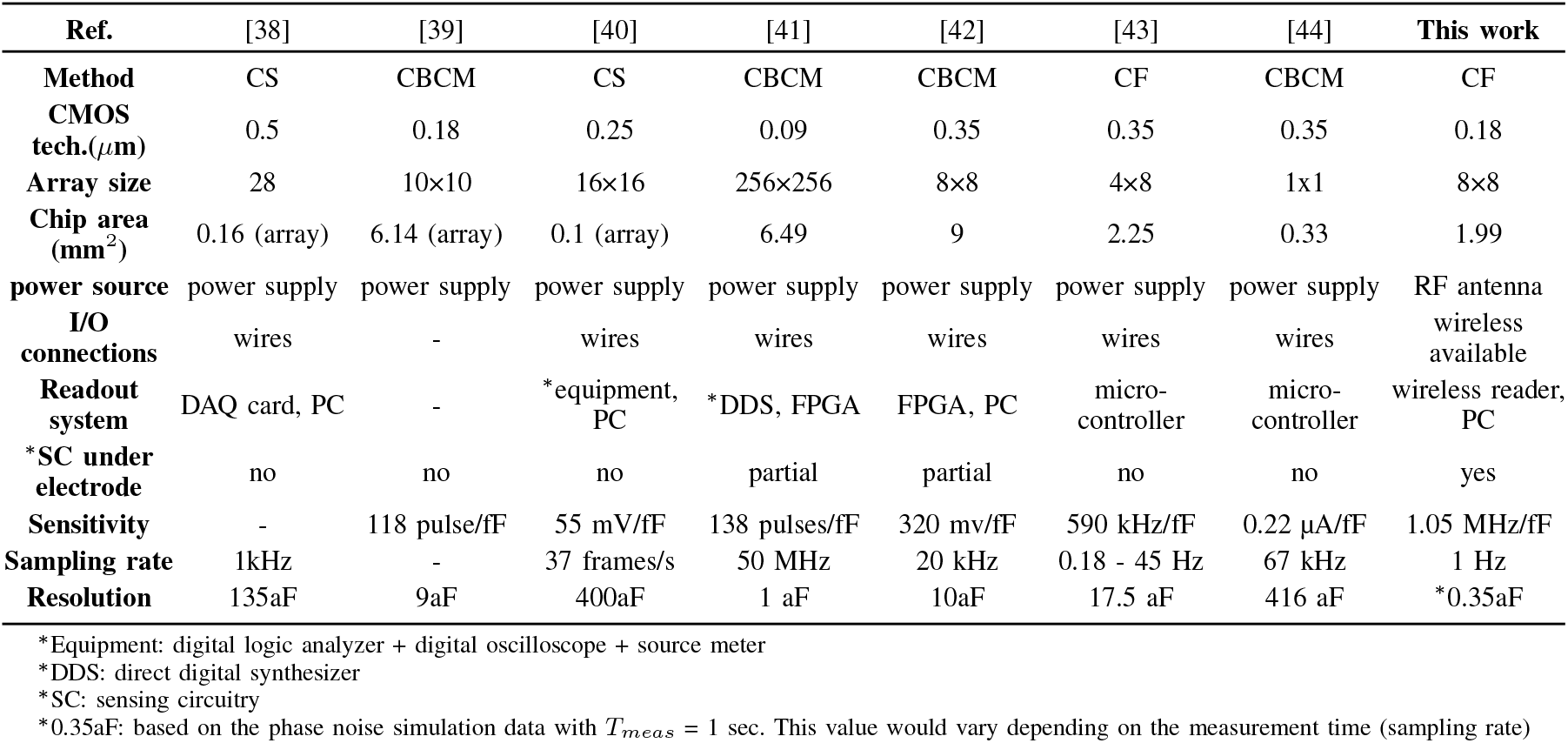
Comparison with the State-of-the-Art.

| Ref. | [38] | [39] | [40] | [41] | [42] | [43] | [44] | This work |
| --- | --- | --- | --- | --- | --- | --- | --- | --- |
| Method | CS | CBCM | CS | CBCM | CBCM | CF | CBCM | CF |
| CMOS tech.($\mu\text{m}$) | 0.5 | 0.18 | 0.25 | 0.09 | 0.35 | 0.35 | 0.35 | 0.18 |
| Array size | 28 | 10×10 | 16×16 | 256×256 | 8×8 | 4×8 | 1×1 | 8×8 |
| Chip area ($\text{mm}^2$) | 0.16 (array) | 6.14 (array) | 0.1 (array) | 6.49 | 9 | 2.25 | 0.33 | 1.99 |
| power source | power supply | power supply | power supply | power supply | power supply | power supply | power supply | RF antenna |
| I/O connections | wires | - | wires | wires | wires | wires | wires | wireless available |
| Readout system | DAQ card, PC | - | *equipment, PC | *DDS, FPGA | FPGA, PC | micro-controller | micro-controller | wireless reader, PC |
| *SC under electrode | no | no | no | partial | partial | no | no | yes |
| Sensitivity | - | 118 pulse/fF | 55 mV/fF | 138 pulses/fF | 320 mv/fF | 590 kHz/fF | 0.22 $\mu\text{A}$/fF | 1.05 MHz/fF |
| Sampling rate | 1kHz | - | 37 frames/s | 50 MHz | 20 kHz | 0.18 - 45 Hz | 67 kHz | 1 Hz |
| Resolution | 135aF | 9aF | 400aF | 1 aF | 10aF | 17.5 aF | 416 aF | *0.35aF |

\*Equipment: digital logic analyzer + digital oscilloscope + source meter
\*DDS: direct digital synthesizer
\*SC: sensing circuitry
\*0.35aF: based on the phase noise simulation data with $T_{meas} = 1$ sec. This value would vary depending on the measurement time (sampling rate)

To our knowledge, the present prototype is the first design that allows a capacitance-sensing lab-on-CMOS chip to power up and transmit sensor data wirelessly using a passive RFID framework. Our design specifically addressed the need for simpler packaging methods in order to facilitate the integration of lab-on-CMOS systems in complex bio-analytical platforms.

Furthermore, we showed that for compact CMOS integration, the sensor array may be implemented using a sensorunder-electrode architecture, allowing all the circuitries necessary for sensing to fit inside a two-loop OCA laid out at the chip’s periphery. In contrast, as reported in Table I, state-of-the-art lab-on-CMOS capacitance sensors typically only partially integrate the sensing circuitry directly under the sensing element.

Our prototype was fabricated in a 0.18 *µ*m commercial CMOS process and tested with a custom-made RFID reader at 915 MHz, *i.e*., in the ISM band range. In our experiments, we successfully powered the chip wirelessly, and we successfully received uplink data from the chip. Furthermore, the sensor data as well as preamble and appendix data markers were demodulated successfully using post-processing, and the demodulated data matched the designed patterns as expected. Moreover, we verified the validity of the senor-under-electrode approach by successfully measuring changes in frequency in relation to air that resulted from several liquid analytes.

The initial work featured herein demonstrates the feasibility of a wireless lab-on-CMOS platform for use in conjuction with a wet test environment. Our prototype was designed for testability. As such, control signals that were not RF-related were provided to the chip externally using I/O pads placed outside the OCA. In a future version, additional circuitry that serves to automatically scan and transmit data from each pixel in the array can be included on-chip thereby obviating the need for I/O pads. Further, although a front-side coupling mode is possible, our work shows that back side coupling the antennas results in fewer issues. The proposed system will facilitate the integration and packaging of a large number of chips in wet environments, paving the way for the inclusion of lab-on-CMOS technology in standard bio-analytical lab practice.

## References

[1] S. Forouhi, R. Dehghani, and E. Ghafar-Zadeh, “CMOS based capacitive sensors for life science applications: A review,” Sensors and Actuators, A: Physical, vol. 297, p. 111531, 2019, doi: 10.1016/j.sna.2019.111531

[2] S. B. Prakash and P. Abshire, “On-Chip Capacitance Sensing for Cell Monitoring Applications,” IEEE Sensors Journal, vol. 7, no. 3, pp. 440–447, 2007, doi: 10.1109/JSEN.2006.889213

[3] B. Senevirathna, S. Lu, M. Dandin, J. Basile, E. Smela, and P. Abshire, “High Resolution Monitoring of Chemotherapeutic Agent Potency in Cancer Cells Using a CMOS Capacitance Biosensor,” Biosensors and Bioelectronics, vol. 142, p. 111501, 2019, doi: 10.1016/j.bios.2019.111501

[4] D. M. Goncalves, R. de Liz, and D. Girard, “Activation of Neutrophils by Nanoparticles,” The Scientific World JOURNAL, vol. 11, pp. 1877–1885, 2011, doi: 10.1100/2011/768350

[5] H. Abdelhamid, T. Z. Salem, M. A. Wahba, D. Mofed, O. E. Morsy, and R. Abdelbaset, “A capacitive sensor for differentiation between virus-infected and uninfected cells,” Sensing and Bio-Sensing Research, vol. 36, p. 100497, 2022, doi: 10.1016/j.sbsr.2022.100497

[6] Y. H. Ghallab and Y. Ismail, “CMOS Based Lab-on-a-Chip: Applications, Challenges and Future Trends,” IEEE Circuits and Systems Magazine, vol. 14, no. 2, pp. 27–47, 2014, doi: 10.1109/MCAS.2014.2314264

[7] J. Abbott, A. Mukherjee, W. Wu, T. Ye, H. S. Jung, K. M. Cheung, R. S. Gertner, M. Basan, D. Ham, and H. Park, “Multi-parametric functional imaging of cell cultures and tissues with a CMOS microelectrode array,” Lab on a Chip, vol. 22, no. 7, pp. 1286–1296, 2022, doi: 10.1039/D1LC00878A

[8] S. Lu, B. Senevirathna, M. Dandin, E. Smela, and P. Abshire, “System Integration of IC Chips for Lab-on-CMOS Applications,” in 2018 IEEE International Symposium on Circuits and Systems (ISCAS). IEEE, 2018, pp. 1–5. ISBN 978-1-5386-4881-0 doi: 10.1109/ISCAS.2018.8351395

[9] M. Dandin, I. Jung, M. Piyasena, J. Gallagher, N. Nelson, M. Urdaneta, C. Artis, P. Abshire, and E. Smela, “Post-CMOS packaging methods for integrated biosensors,” in Proceedings of IEEE Sensors, 2009. ISBN 9781424445486 doi: 10.1109/ICSENS.2009.5398540

[10] T. Datta-Chaudhuri, E. Smela, and P. A. Abshire, “System-on-Chip Considerations for Heterogeneous Integration of CMOS and Fluidic Bio-Interfaces,” IEEE Transactions on Biomedical Circuits and Systems, vol. 10, no. 6, pp. 1129–1142, 2016, doi: 10.1109/TBCAS.2016.2522402

[11] L. Lantz and M. Pecht, “Ion transport in encapsulants used in microcircuit packaging,” IEEE Transactions on Components and Packaging Technologies, vol. 26, no. 1, pp. 199–205, 2003, doi: 10.1109/TCAPT.2002.806183

[12] Y. Huang and A. J. Mason, “Lab-on-CMOS: Integrating Microfluidics and Electrochemical Sensor on CMOS,” in 2011 6th IEEE International Conference on Nano/Micro Engineered and Molecular Systems. IEEE, 2011, pp. 690–693. ISBN 978-1-61284-775-7 doi: 10.1109/NEMS.2011.6017448

[13] T. Datta-Chaudhuri, P. Abshire, and E. Smela, “Packaging Commercial CMOS Chips for Lab on a Chip Integration,” Lab on a Chip, vol. 14, no. 10, p. 1753, 2014, doi: 10.1039/c4lc00135d

[14] H. Yin, L. Li, and A. J. Mason, “Screen-Printed Planar Metallization for Lab-on-CMOS with Epoxy Carrier,” in 2016 IEEE International Symposium on Circuits and Systems (ISCAS). IEEE, 2016, pp. 2887–2890. ISBN 978-1-4799-5341-7 doi: 10.1109/ISCAS.2016.7539196

[15] L. Li, H. Yin, and A. J. Mason, “Epoxy Chip-in-Carrier Integration and Screen-Printed Metalization for Multichannel Microfluidic Lab-on-CMOS Microsystems,” IEEE Transactions on Biomedical Circuits and Systems, vol. 12, no. 2, pp. 416–425, 2018, doi: 10.1109/TB-CAS.2018.2797063

[16] Jae-Ryong Cha and Jae-Hyun Kim, “Novel Anti-collision Algorithms for Fast Object Identification in RFID System,” in 11th International Conference on Parallel and Distributed Systems (ICPADS’05). IEEE, pp. 63–67. ISBN 0-7695-2281-5 doi: 10.1109/ICPADS.2005.204

[17] Weilian Su, N. Alchazidis, and T. Ha, “Multiple RFID Tags Access Algorithm,” IEEE Transactions on Mobile Computing, vol. 9, no. 2, pp. 174–187, 2010, doi: 10.1109/TMC.2009.106

[18] H. Vogt, “Multiple Object Identification with Passive RFID Tags,” in IEEE International Conference on Systems, Man and Cybernetics. IEEE, 2003, p. 6. ISBN 0-7803-7437-1 doi: 10.1109/IC-SMC.2002.1176119

[19] Y. Gilpin, C.-Y. Lin, and M. P. Dandin, “Evaporation and focus degradation mitigation in in-incubator live cell imaging for capacitance lab-on-cmos microsystem calibration,” Review of Scientific Instruments, vol. 96, no. 6, 2025, doi: 10.1063/5.0271101

[20] C.-Y. Lin and M. P. Dandin, “Machine learning identification and classification of mitosis and migration of cancer cells in a lab-on-cmos capacitance sensing platform,” IEEE Journal of Biomedical and Health Informatics, vol. 29, no. 2, pp. 1504–1513, 2024, doi: 10.1109/JBHI.2024.3486251

[21] K. Smith, C.-Y. Lin, Y. Gilpin, E. Wayne, and M. P. Dandin, “Measuring and modeling macrophage proliferation in a lab-on-cmos capacitance sensing microsystem,” Frontiers in Bioengineering and Biotechnology, vol. 11, p. 1159004, 2023, doi: 10.3389/fbioe.2023.1159004

[22] B. Senevirathna, S. Lu, M. P. Dandin, E. Smela, and P. Abshire, “Correlation of capacitance and microscopy measurements using image processing for a lab-on-cmos microsystem,” IEEE Transactions on Biomedical Circuits and Systems, vol. 13, no. 6, pp. 1214–1225, 2019, doi: 10.1109/TBCAS.2019.2926836

[23] B. Senevirathna, S. Lu, M. P. Dandin, J. Basile, E. Smela, and P. Abshire, “Real-time measurements of cell proliferation using a lab-on-cmos capacitance sensor array,” IEEE Transactions on Biomedical Circuits and Systems, vol. 12, no. 3, pp. 510–520, 2018, doi: 10.1109/TB-CAS.2018.2821060

[24] A. Hajimiri, S. Limotyrakis, and T. Lee, “Jitter and Phase Noise in Ring Oscillators,” IEEE Journal of Solid-State Circuits, vol. 34, no. 6, pp. 790–804, 1999, doi: 10.1109/4.766813

[25] S. Suman, K. G. Sharma, and P. K. Ghosh, “Analysis and Design of Current Starved Ring VCO,” in 2016 International Conference on Electrical, Electronics, and Optimization Techniques (ICEEOT). IEEE, 2016, pp. 3222–3227. ISBN 978-1-4673-9939-5 doi: 10.1109/ICEEOT.2016.7755299

[26] S. Docking and M. Sachdev, “A Method to Derive an Equation for the Oscillation Frequency of a Ring Oscillator,” IEEE Transactions on Circuits and Systems I: Fundamental Theory and Applications, vol. 50, no. 2, pp. 259–264, 2003, doi: 10.1109/TCSI.2002.808235

[27] K. Finkenzeller and D. Muller, RFID Handbook: Fundamentals and Applications in Contactless Smart Cards, Radio Frequency Identification and Near-Field Communication. Wiley Telecom, 2010.

[28] L. R. Carley, G. Colak, L. Chomas, L. Pileggi, and K. Mai, “Technologies for Secure RFID Authentication of Medicinal Pills and Capsules,” in 2016 IEEE International Conference on RFID Technology and Applications (RFID-TA). IEEE, 2016, pp. 10–15. ISBN 978-1-5090-1408-8 doi: 10.1109/RFID-TA.2016.7750745

[29] T. Delbruck, I. A. M. Elfadel, S. Muzaffar, G. Haessig, B. Wang, Bermak, R. Graca, L. Camunas-Mesa, B. Senevirathna, P. Abshire, Linares-Barranco, S. Afshar, S.-C. Liu, R. M. Wang, P. Dudek, S. Carey, J. de la Rosa, M. Dandin, S. Lu, V. Frick, T. Serrano-Gotarredona, P. Lopez, M. Payvand, A. Madhavan, E. Fossum, J. C. V. Tieck, I. Williams, Y. Liu, T. Constandinou, A. Serb, R. Carmona-Galan, R. Nawrocki, and W. D. Leon-Salas, “Lessons Learned the Hard Way,” in 2020 IEEE International Symposium on Circuits and Systems (ISCAS). IEEE, 2020, pp. 1–18. ISBN 978-1-7281-3320-1 doi: 10.1109/ISCAS45731.2020.9180983

[30] Koji Kotani and Takashi Ito, “High Efficiency CMOS Rectifier Circuit with Self-Vth-Cancellation and Power Regulation Functions for UHF RFIDs,” in 2007 IEEE Asian Solid-State Circuits Conference. IEEE, 2007, pp. 119–122. ISBN 978-1-4244-1359-1 doi: 10.1109/ASSCC.2007.4425746

[31] A. S. Bakhtiar, M. S. Jalali, and S. Mirabbasi, “A High-Efficiency CMOS Rectifier for Low-Power RFID Tags,” in 2010 IEEE International Conference on RFID (IEEE RFID 2010). IEEE, 2010, pp. 83–88. ISBN 978-1-4244-5742-7 doi: 10.1109/RFID.2010.5467271

[32] D. Ash, “A Comparison Between OOK/ASK and FSK Modulation Techniques for Radio Links,” Computer Science, 2002.

[33] D. De Donno, F. Ricciato, and L. Tarricone, “Listening to Tags: Uplink RFID Measurements With an Open-Source Software-Defined Radio Tool,” IEEE Transactions on Instrumentation and Measurement, vol. 62, no. 1, pp. 109–118, 2013, doi: 10.1109/TIM.2012.2212513

[34] C. Yue and S. Wong, “On-chip Spiral Inductors with Patterned Ground Shields for Si-based RF ICs,” IEEE Journal of Solid-State Circuits, vol. 33, no. 5, pp. 743–752, 1998, doi: 10.1109/4.668989

[35] K. Murata, T. Hosaka, and Y. Sugimoto, “Effect of a Ground Shield of a Silicon on-Chip Spiral Inductor,” in 2000 Asia-Pacific Microwave Conference. Proceedings (Cat. No.00TH8522). IEEE, 2000, pp. 177–180. ISBN 0-7803-6435-X doi: 10.1109/APMC.2000.925754

[36] N. Talwalkar, C. Yue, and S. Wong, “Analysis and Synthesis of On-Chip Spiral Inductors,” IEEE Transactions on Electron Devices, vol. 52, no. 2, pp. 176–182, 2005, doi: 10.1109/TED.2004.842535

[37] G. Benelli and A. Pozzebo, “RFID Under Water: Technical Issues and Applications,” in Radio Frequency Identification from System to Applications. InTech, 2013, doi: 10.5772/53934

[38] S. B. Prakash and P. Abshire, “Tracking Cancer Cell Proliferation on a CMOS Capacitance Sensor Chip,” Biosensors and Bioelectronics, vol. 23, no. 10, pp. 1449–1457, 2008, doi: 10.1016/j.bios.2007.12.015

[39] S. Forouhi, R. Dehghani, and E. Ghafar-Zadeh, “High Throughput Core-CBCM CMOS Capacitive Sensor for Life Science Applications,” in 2018 IEEE Canadian Conference on Electrical & Computer Engineering (CCECE). IEEE, 2018, pp. 1–4. ISBN 978-1-5386-2410-4 doi: 10.1109/CCECE.2018.8447694

[40] N. Couniot, L. A. Francis, and D. Flandre, “A 16 x 16 CMOS Capacitive Biosensor Array Towards Detection of Single Bacterial Cell,” IEEE Transactions on Biomedical Circuits and Systems, vol. 10, no. 2, pp. 364–374, 2016, doi: 10.1109/TBCAS.2015.2416372

[41] F. Widdershoven, A. Cossettini, C. Laborde, A. Bandiziol, P. P. van Swinderen, S. G. Lemay, and L. Selmi, “A CMOS Pixelated Nanocapacitor Biosensor Platform for High-Frequency Impedance Spectroscopy and Imaging,” IEEE Transactions on Biomedical Circuits and Systems, vol. 12, no. 6, pp. 1369–1382, 2018, doi: 10.1109/TBCAS.2018.2861558

[42] G. Nabovati, E. Ghafar-Zadeh, A. Letourneau, and M. Sawan, “Smart Cell Culture Monitoring and Drug Test Platform Using CMOS Capacitive Sensor Array,” IEEE Transactions on Biomedical Engineering, vol. 66, no. 4, pp. 1094–1104, 2019, doi: 10.1109/TBME.2018.2866830

[43] B. Senevirathna, S. Lu, N. Renegar, M. Dandin, E. Smela, and P. Abshire, “System on a Chip for Automated Cell Assays using a Lab-on-CMOS Platform,” in 2019 IEEE International Symposium on Circuits and Systems (ISCAS). IEEE, 2019, pp. 1–5. ISBN 978-1-7281-0397-6 doi: 10.1109/ISCAS.2019.8702702

[44] H. O. Tabrizi, O. Farhanieh, Q. Owen, S. Magierowski, and E. Ghafar-Zadeh, “Wide Input Dynamic Range Fully Integrated Capacitive Sensor for Life Science Applications,” IEEE Transactions on Biomedical Circuits and Systems, vol. 15, no. 2, pp. 339–350, 2021, doi: 10.1109/TB-CAS.2021.3075348

